# Multi-Modal Toxicological Evaluation of NiO nanoparticles and ionic nickel in a Human Lung-Cardiac Co-Culture System

**DOI:** 10.64898/2026.07.10.737738

**Authors:** Wesam Darwish, Sophie Kussauer, Mohammad Almasaleekh, Sebastiano Di Bucchianico, Ralf Zimmermann, Robert David

## Abstract

Nickel is a widespread environmental and occupational contaminant associated with respiratory and cardiovascular toxicity, yet the mechanisms linking pulmonary exposure to adverse cardiac effects remain poorly understood. This study aimed to establish and evaluate a human in vitro lung–heart co-culture model for investigating cardiovascular responses following pulmonary exposure. Human alveolar epithelial A549 cells were exposed at the air–liquid interface to different concentrations of NiO nanoparticles or NiCl_2_ for 4- and 24-hours. Following cloud exposure, A549 cells were co-cultured with human induced pluripotent stem cell-derived cardiomyocytes (hiPSC-CMs). Cytotoxicity, metabolic activity, cytokine release, DNA damage, epigenetic alterations, and cardiac electrophysiological function were assessed. Nickel translocation across the epithelial barrier was quantified to facilitate interpretation of downstream cardiomyocyte effects. Exposure to both nickel forms induced cytotoxicity and resulted in measurable nickel translocation into the basolateral compartment. NiCl_2_ exhibited a time-dependent increase in basolateral nickel concentrations, whereas NiO translocation remained relatively stable over time. Cytokine profiling revealed selective induction of IL-8 and IL-18, with no significant changes in IL-1β, IL-6, IL-10, or TNF-α. Genotoxicity analyses demonstrated cell type-specific responses, characterized by delayed DNA strand breaks in A549 cells and early but transient DNA damage in hiPSC-CMs. Oxidative DNA damage was particularly pronounced in hiPSC-CMs following NiCl_2_ exposure. Global DNA methylation was reduced in hiPSC-CMs without corresponding changes in DNA methyltransferase activity. Electrophysiological assessment showed transient increases in conduction velocity, while beating frequency and field potential duration remained largely unaffected. Overall, the lung–heart co-culture model successfully captured both pulmonary and cardiac responses to nickel exposure and provided evidence for direct and indirect mechanisms of cardiotoxicity. Nickel translocation across the epithelial barrier, together with inflammatory and oxidative stress-related signalling, may contribute to downstream cardiac effects. These findings highlight the utility of this human-relevant platform for investigating systemic cardiovascular consequences of inhaled toxicants.

## Introduction

Nickel is a widespread environmental and occupational contaminant to which humans are routinely exposed. Natural sources, such as volcanic activity, wildfires, and weathering, contribute to background environmental levels, whereas industrial processes, including metal refining, welding, and fuel combustion, represent major occupational exposures (Rizwan et al., 2024). Additional exposure occurs through tobacco smoke and dietary intake, particularly from plant-derived foods and seafood (Genchi et al., 2020). Owing to its well-established toxic and carcinogenic properties, regulatory agencies have implemented strict exposure limits, including a tolerable daily intake of 13 µg kg^-1^ body weight (EFSA, 2020), and an occupational exposure limit of 0.01 mg m^-^³ for inhalable nickel compounds (SCOEL, 2011).

Nickel and its compounds exhibit substantial differences in toxicological behaviour and carcinogenic potential. Nickel compounds are classified as Group 1 carcinogens, whereas metallic nickel is classified as Group 2B by the International Agency for Research on Cancer (IARC, 1990). Notably, poorly soluble nickel species generally display greater carcinogenicity than soluble forms, a difference attributed to variations in cellular uptake, intracellular persistence, and interference with cell cycle regulation (Ouyang et al., 2009). Chronic nickel exposure has been associated with a range of respiratory disorders, including asthma (Hong et al., 1986; Gül et al., 2007), pulmonary fibrosis (Berge et al., 2003; Cao et al., 2024), lung cancer, and chronic obstructive pulmonary disease (Albayrak et al., 2023). Beyond its local pulmonary effects, accumulating evidence suggests that inhaled nickel can translocate from the lungs to the cardiovascular system (Wallenborn et al., 2007; Nakane, 2012) and contribute to adverse cardiac outcomes, including cardiac dysfunction and an increased risk of congenital heart disease (Lippmann et al., 2006; Zhang et al., 2019; Cheek et al., 2024).

At the cellular level, insoluble nickel nanoparticles are internalized by lung cells primarily through particle uptake mechanisms, including phagocytic and endocytic pathways (Zambelli et al., 2016; Di Bucchianico et al., 2018). In contrast, ionic nickel enters cells as dissolved Ni²⁺ ions through ion channel- and transporter-mediated mechanisms (Refsvik et al., 1995). Once internalized, nickel can induce oxidative stress, inflammation, and alterations in gene expression, lipid metabolism, and apoptotic pathways (Lü et al., 2009; Fidan et al., 2024). Importantly, the solubility of nickel compounds strongly influences their biological fate: soluble forms readily enter systemic circulation, whereas particulate or poorly soluble species tend to persist in pulmonary tissues, potentially acting as a sustained source of metal ions (Schaumlöffel, 2012).

Because inhaled toxicants can influence distant organs through biological mediators and chemical and/or particles translocation (Gorr et al., 2015; Elder et al., 2006), lung-heart interactions are potentially a key determinant of nickel-induced systemic toxicity. Pulmonary exposure to nickel in vivo can depress cardiac oxygen consumption, increase mitochondrial ROS production (Garcés et al., 2021), and disrupt calcium signalling (Hobai et al., 2000). These effects may potentially lead to arrhythmia (Lou et al., 2013) and exacerbated case of atherosclerosis (Kang et al., 2011), even in the absence of direct particle translocation (Hobai et al., 2000; Kang et al., 2011; Garcés et al., 2021).

Despite growing recognition of nickel-induced systemic toxicity, the mechanisms underlying lung-to-heart communication following inhalation exposure remain poorly understood. Most existing studies rely either on isolated lung cell models or on in vivo approaches, neither of which adequately captures dynamic inter-organ signalling in a human-relevant context. Although conditioned-medium studies have demonstrated that factors released from nickel-exposed lung cells can impair cardiomyocyte function (Gorr et al., 2015), these approaches lack physiologically relevant exposure conditions and do not reflect the dynamic temporal interactions that occur between organs. Consequently, there is a critical need for advanced in vitro models that enable direct investigation of pulmonary-cardiac interactions under realistic exposure scenarios. Such models would also facilitate functional assessment of cardiomyocyte electrophysiology, enabling the detection of arrhythmogenic and contractility-related toxic effects.

Air-liquid interface (ALI) systems provide a physiologically relevant platform for inhalation toxicology by allowing the direct deposition of aerosols onto lung epithelial cells, thereby closely mimicking respiratory exposure. When combined with human induced pluripotent stem cell-derived cardiomyocytes (hiPSC-CMs), these systems offer a unique opportunity to investigate secondary cardiac effects resulting from pulmonary exposure (Pourrier et al., 2020; Ng et al., 2022). This integrated approach addresses key limitations of traditional animal models, including interspecies variability, ethical concerns, and the substantial time and cost associated with in vivo studies (Hartung, 2024). In addition, it enables functional assessment of cardiomyocyte electrophysiology and contractility, providing sensitive endpoints for the detection of cardiotoxic effects.

The aim of this study was to establish and validate a human-relevant ALI lung-heart co-culture model comprising A549 lung epithelial cells and hiPSC-derived cardiomyocytes to investigate both pulmonary and secondary cardiac responses to nickel exposure. Specifically, we compared nickel oxide nanoparticles (NiO-NPs), representing a poorly soluble particulate form of nickel, with nickel chloride (NiCl_2_), a soluble ionic form, to determine how physicochemical properties influence biological responses. Using this platform, we evaluated cytotoxicity, inflammatory signalling, DNA damage, and epigenetic modifications, and examined how these responses are propagated to cardiomyocytes to affect electrophysiological function. By integrating multi-endpoint analyses with a physiologically relevant exposure system, this study provides mechanistic insight into lung-heart communication following nickel inhalation and establishes a versatile human-relevant platform for evaluating the cardiotoxic potential of airborne contaminants.

## Materials and Methods

### Cardiac differentiation of hiPSCs

For cardiac differentiation human iPSC wild-type cells (HDF, SC950A-1) were used. After thawing the iPSC cells were cultured under hypoxic conditions at 37 °C in iPS-Brew XF medium (Miltenyi Biotec, 130-104-368). Culture flasks were coated with biolaminin-521 (Biolamina, KN521-05) overnight at 2-8 °C using a 5 µg ml^-1^ concentration in PBS. Cardiac differentiation to iPSCs-CM was performed as described before [Skorska et l. 2022]. Passaging and seeding of the cells for differentiation were both performed using the Accutase (Sigma-Aldrich, A6964-500) and iPS-Brew supplemented with 5 µM ROCK inhibitor (StemMACs Y27632, Miltenyi Biotec, 130-103-922), for 24 h.

To start the cardiac differentiation, cells were seeded onto biolaminin 521-coated well plates and cultured under normoxic conditions in cardiac differentiation medium consisting of RPMI 1640 (+Glutamax; Gibco, 61870-010), with 2 % B27 supplement without insulin (Gibco, A1895601), 1 % sodium pyruvat (Gibco, 11360070), 200 µM ascorbic acid (Merck, A8960- 5g) and 1 % ZellShield (Minerva Biolabs, 13-0050). In order to induce mesoderm, the medium was supplemented with the following substances for two days: 1 µM Chir99021 (Miltenyi Biotech, 130-103-926), 5 ng ml^-1^ bFGF (Miltenyi Biotec, 130-104-918), 5 ng ml^-1^ BMP4 (Miltenyi Biotec, 130-111-168) and 9 ng ml^-1^ ActivinA (Miltenyi Biotec, 130-115-012). The following cardiac induction was driven by a supplementation of cardiac differentiation medium using B27 supplement with insulin (Gibco, A3695201) and 5 mM IWP2 (Tocris, 3533) for a period of seven days. Thereafter, the cells were cultivated for further three days in the absence of IWP2 supplementation. To perform a metabolic selection, the cultivation of cells was conducted in a selection medium composed of RPMI 1640 without glucose (PAN Biotech, P04-17545), 100 µM β-mercaptoethanol (Sigma-Aldrich, M3148), 1 % ZellShield and 2,3 mM sodium lactate (Sigma-Aldrich, L7900-100ml) for four days. This was followed by a subsequent three-day period in an insulin-containing cardiac differentiation medium.

For co-culture experiments, the culture surfaces for the hiPSC-CM were coated with biolaminin-521 (12-well plate) or Matrigel (1:50 Corning, 356234; glass surface of MEA) and the CMs were seeded with 400,000 cells for each well inside 12-well plate or one microelectrode array.

### A549 cell culture

The human alveolar epithelial A549 cells (ATCC, CCL-185) were thawed in DMEM/F-12 GlutaMAX medium (Gibco, 10565018) supplemented with 1 % (v/v) Penicillin-Streptomycin (Gibco, 15140122) and 10 % (v/v) heat-inactivated fetal bovine serum (FBS; Sigma-Aldrich, F7524) and maintained at 37 °C and 5 % CO₂. After the first passage, the FBS concentration in the medium was reduced to 5 %, and the medium is hereafter referred to as A549 medium. Cells were subsequently sub-cultured three times before ALI exposure.

For co-culture experiments, A549 cells were seeded onto transferrable Transwell inserts at 1×10⁵ cells/cm², using 12-well inserts (CellQart, 9310412) for cytotoxicity and genetic-related analyses and 24-well inserts (CellQart, 9320412) for cardiomyocyte functional studies. The basolateral compartment of each 12-transwell insert was filled with 1.5 mL of DMEM/F-12 supplemented with 1 % Penicillin-Streptomycin and 5 % FBS. After 24 hours of incubation, the apical medium was aspirated to establish ALI, and the basolateral medium was replaced with cardiac differentiation medium with insulin.

### Assessment of Media and Cell Compatibility for Establishing A549-hiPSC-CM Co-Culture System

To evaluate the compatibility of lung epithelial A549 cells with hiPSC-CMs for co-culture experiments, both media and cell compatibility were assessed.

For media compatibility, three media conditions were tested: (i) DMEM/F-12 medium supplemented with 1 % Penicillin-Streptomycin and 5 % fetal bovine serum (hereafter referred to as “A549 medium”); (ii) RPMI medium supplemented with B27 supplement with insulin and 1 % sodium pyruvate and 200 mM ascorbic acid and 1 % ZellShield (hereafter referred to as “cardiac differentiation medium” or “hiPSC-CM medium”; and (iii) a 1:1 (v/v) mixture of the two media. A total of 45,000 A549 cells cm^-2^ were seeded into 12-well Transwell insert, and 300,000 hiPSC-CMs cm^-2^ were seeded into each well of a 6-well microelectrode array (MEA; Multi-Channel Systems, 60-6wellMEA200/30iR-Ti-rcr) pre-coated with Matrigel. Field potential properties of hiPSC-CMs were recorded prior to medium application and subsequently at 4, 24, 55, and 73 hours using an MEA2100 system and Cardio2D software (Multi-Channel Systems, Reutlingen). Cytotoxicity and metabolic activity were assessed for both A549 cells and hiPSC-CMs under each medium condition using Cytotoxicity Detection Kit (Roche, 4744926001) and CellTiter-Blue® reagent (Promega, G8081).

To evaluate compatibility between cell types in co-culture, 90,000 A549 cells/cm^-2^ were seeded into 24-well Transwell inserts and cultured either alone or placed onto one well MEAs seeded with 400,000 hiPSC-CMs. The A549 insert was attached to the MEA using a customized 3D-printed lid (Mau et al., 2021). Cytotoxicity and metabolic activity were measured in monocultures of A549 cells, monocultures of hiPSC-CMs, and co-cultures of both cell types using the aforementioned assays. Effects on cardiac field potentials were assessed at 0, 4 and 24 hours of co-culture using the mentioned MEA technology.

Further methodological details regarding cytotoxicity, metabolic activity, and microelectrode array analysis are provided in the corresponding sections below.

### Air-Liquid Interface Exposure and Co-culture

The day after establishing the ALI condition, the Transwell inserts were exposed to either nickel oxide nanoparticles (NiO; ChemPur, CFF-NP032) or their ionic equivalent, nickel chloride (NiCl_2_; Sigma-Aldrich, 223387), using the VITROCELL Cloud Alpha 6 exposure system. Three concentrations of the nebulization suspensions were freshly prepared for each compound in ultrapure water spiked with 1 % (v/v) HBSS containing calcium and magnesium (Gibco; 14025-050), corresponding to 250, 500, and 750 µg mL^-1^ based on the nickel content. For nebulization, 200 µL of the prepared suspensions was aerosolized using a 3–5 µm nebulizer, and the generated aerosol cloud was allowed to settle over the inserts for 5 minutes to ensure uniform deposition. Two control conditions were included in all experiments: an incubator control (IC), in which the cells were left untreated at ALI, and a solvent control (SC), in which cells were exposed to ultrapure water supplemented with 1 % HBSS containing calcium and magnesium mimicking the treatment conditions.

Following each nebulization, the co-cultures were assembled differently depending on the downstream analysis. For MEA analysis, exposed A549 cells in 24-well inserts were positioned on top of 300,000 hiPSC-CM using a 3D-printed lid, which had been seeded on 1-well microelectrode array (MEA) chips (60MEA100/10iR-Ti; 60MEA200/30iR-Ti, Multi-Channel Systems, Reutlingen). The co-culture was maintained using fresh cardiac differentiation medium. Following the recording of baseline field potential measurements, the spent medium from the inserts was collected, mixed in a 1:1 ratio with fresh insulin-containing cardiac differentiation medium. The resultant mixture was added back to the hiPSC-CM on the MEA arrays to continue the co-culture for the subsequent endpoints. For all other assays, including cytotoxicity, IL-8 secretion, DNA damage, and epigenetic studies, the exposed A549 inserts were placed into 12-well plates seeded with 400,000 hiPSC-CMs. In all cases, co-cultures were maintained in a humidified incubator, and analyses were performed at 0, 4, and 24hours post-exposure.

Published data indicate that the VITROCELL® Cloud Alpha 6 system achieves a stable aerosol deposition efficiency of 15.6 % ± 0.6 % (Zimmermann et al., 2023). Accordingly, the estimated deposition doses for nickel oxide and nickel ions are 280, 560, and 840 ng cm^-^² for nebulization solutions of 250, 500, and 750 µg mL^-1^, respectively. All results are interpreted based on estimated deposited doses (ng Ni cm^-^²), while nebulization concentrations are provided for preparation reference.

### Atomic Absorption Spectroscopy

The nickel content in basolateral medium samples was quantified using an Atomic Absorption Spectrometer (contrAA 800 D HR-CS atomic absorption spectrometer, Analytik Jena AG, Germany) equipped with a Xenon short arc lamp and a graphite furnace atomizer. Method parameters were defined in the instrument software ASpect CS 2.3.1.0. Nr.: 0002: analytical wavelength 232.0030 nm and IBC.

Calibration was performed daily using a certified nickel AAS standard solution (ROTI®Star, 1000 mg L−Ni in 2 % HNO_3_) at concentrations of 2, 4, 6, 8, and 10 μg L^-1^ for the lower concentration range and 6, 12, 18, 24, and 30 μg L^-1^ for the higher concentration range. The standard solutions were prepared by the instrument itself using the highest standard solution (10 or 30 μg L^-1^), with 0.5% HNO₃ serving as the diluent and as a blank. The 0.5% HNO₃ solution was prepared using 69 % nitric acid, supra pur, Carl Roth GmbH and Milli-Q-water. Control standards were prepared by diluting a VWR® AVS Titrinorm, Nickel standard solution 1 g L^-1^ to 5 respectively 20 μg L^-1^. Every measured solution was added by 5 µl matrixmodifier 10 g L^-1^ Mg(NO_3_)_2_ (99,999 %) in H_2_O (Roti®Star) to ensure high pyrolyses temperatures. The calibration curve was established with a 98.50% confidence interval to ensure reliable quantification of nickel. Pre-dilutions with 0.5% HNO₃ were often necessary to bring the nickel concentration within the calibration range. Eppendorf pipettes and corresponding tips were used for all liquid handling to ensure precision and accuracy. Each sample and each standard solution was measured three times to verify reproducibility in quantifying nickel concentrations. Samples were stored at -80°C until analysis.

### Cytotoxicity and metabolic activity assays

The cytotoxicity of the monocultures and co-culture was assessed using the lactate dehydrogenase (LDH)-based Cytotoxicity Detection Kit (Roche, 4744926001) following the manufacturer’s instructions. The apical compartments were washed with 0.5 mL of prewarmed HBSS (Gibco, 14025-050), and the resulting suspension of the apical wash was collected along with the basolateral medium for analysis. An insert of untreated A549 cells incubated with 5 % (v/v) lysis solution for 20 minutes served as a positive control. Collected samples were centrifuged at 250 ×g for 5 minutes, and 50 µL of each supernatant was transferred into a 96-well plate. The LDH reaction was initiated by adding 50 µL of the kit’s reaction mixture to each well. Plates were incubated in the dark for 20 minutes, and absorbance was measured at 492 nm and 620 nm using a microplate reader. The absorbances of the apical wash and basolateral medium reactions were summed for each co-culture condition, and cytotoxicity percentages were calculated relative to the positive control.

For metabolic activity, 0.5 mL of fresh cardiomyocyte medium containing 10 % (v/v) CellTiter-Blue® reagent (Promega, G8081) was added to both the apical and basolateral compartments after the previously mentioned washing step. Plates were incubated for 1 hour at 37 °C in a 5% CO₂ humidified incubator. Fluorescence was measured at 560Ex/590Em for the apical and basolateral compartments separately, and the signals were summed to calculate the metabolic activity of the co-culture. The summed values from the apical and basolateral compartments in both assays represent the overall toxic impact on the co-culture system.

### Cytokines Quantification

The concentrations of five pro-inflammatory cytokines (IL-1β, IL-6, IL-8, IL-18 and TNF-α) and the anti-inflammatory cytokine (IL-10) were quantified using both colorimetric and luminescence-based immunoassays according to the manufacturers’ instructions.

Levels of IL-8 and IL-18 were determined using DuoSet ELISA kits (R&D Systems, DY208 and DY318). Briefly, 96-well plates were coated overnight at room temperature with 100 µL of capture antibody. Plates were then washed three times with wash buffer and blocked for 1 hour with 300 µL of blocking buffer. Following an additional wash step, 100 µL of basolateral medium or standards were added and incubated for 2 hours. Standard curves were generated using serial two-fold dilutions, ranging from 31.2 to 2000 pg mL^-1^ for IL-8 and from 11.7 to 750 pg mL^-1^ for IL-18. After washing, 100 µL of detection antibody was added and incubated for 2 hours, followed by incubation with streptavidin-horseradish peroxidase for 20 minutes. Plates were washed again before adding 100 µL of 3,3′,5,5′-tetramethylbenzidine (TMB) substrate and incubated for 20 minutes. The reaction was stopped with 50 µL of 2 N H₂SO₄, and absorbance was measured at 450 nm with a reference wavelength of 570 nm.

Additional cytokines, including IL-1β, IL-6, IL-10, and TNF-α, were quantified using luminescence-based ELISA kits (Promega; W6010, W6030, W6070, W6050). For these assays, 50 µL of basolateral medium were added in duplicate to white 96-well plates together with seven-point standard curves prepared by serial dilution (18.2–25,000 pg mL^-1^). Subsequently, 50 µL of antibody mixture was added, and plates were briefly mixed before incubation at 37 °C in a CO₂ incubator for 60 minutes. After equilibration to room temperature, 25 µL of luminescence detection reagent was added. Plates were shaken briefly and incubated for 4 minutes prior to luminescence measurement using a multimode microplate reader.

For IL-1β, assay conditions were adjusted according to the manufacturer’s recommendations. Standard curves ranged from 21.7 to 40,000 pg mL^-1^ using 3.5-fold serial dilutions, and incubation time was extended to 75 minutes.

### DNA Extraction and Quantification

The co-culture model was exposed to nebulization solutions of 0 and 750 µg mL^-1^ NiO or NiCl_2_ based on nickel content, then incubated for 24 hours inside a humified incubator. DNA extraction was performed using the FitAmp Blood and Cultured Cell DNA Extraction Kit (EpigenTek, P-1018) according to the manufacturer’s instructions. Briefly, A549 cells from 10 Transwell inserts (12-well format) and their corresponding cardiomyocytes from the basolateral compartment were pooled separately. Cell pellets were obtained by centrifugation at 2000 rpm for 3 minutes, and 1 × 10⁶ cells were resuspended in 200 µL of suspending buffer. After vortexing, 4 µL of DNA digestion reagent was added, followed by another brief vortex step and incubation at 65 °C for 15 minutes. Next, 300 µL of DNA capture buffer was added, and the mixture was transferred to a spin column and centrifuged at 12,000 rpm for 45 seconds. The flow-through was discarded, 300 µL of 70 % ethanol (Carl Roth, 1HPH.1) was then added, and samples were centrifuged at the same speed for 30 seconds. After discarding the flow-through, 200 µL of 90 % ethanol was added and centrifuged again, followed by a second wash with 90 % ethanol and a prolonged centrifugation step of 40 seconds. DNA was eluted into fresh tubes using 15 µL of elution buffer and collected by centrifugation for 20 seconds.

DNA quantification was performed using the QuantiFluor® ONE dsDNA System (Promega, E4870). Following the manufacturer’s recommendations, 1 µL of extracted DNA was mixed with 200 µL of the supplied dye in a black 96-well plate. Seven standards ranging from 0.2 ng µL^-1^ to 400 ng µL^-1^ were included in each assay. Samples were homogenized by shaking at 600 rpm for 15 seconds and incubated for 5 minutes protected from light. Fluorescence was measured at 504Ex/531Em, and DNA quantities were calculated using the corresponding standard curve.

### 5-Methylcytosine Quantification

The levels of 5-methylcytosine in DNA from exposed cells were assessed using MethylFlash Methylated DNA 5-mC Quantification Fluorometric Kit (EpigenTek, P-1035). The assay plate was first coated with 80 µL of binding solution, after which 100 ng of sample DNA was added to the designated wells. In parallel, 1 µL of the negative control and 1 µL of the positive control containing 50% 5-methylcytosine were added. The plate was sealed and incubated at 37 °C for 90 minutes. The wells were then emptied and washed three times with 150 µL of 1× wash buffer. Next, 50 µL of 1 µg mL^-1^ capture antibody was added, followed by incubation for 60 minutes at room temperature. After three additional washes, 50 µL of 200 ng mL^-1^ detection antibody was added and the plate was incubated for 30 minutes. The plate was washed four times, and 50 µL of enhancer solution was added and incubated for 30 minutes. This was followed by five washes with diluted wash buffer and a final wash with 150 µL of DPBS. Finally, 50 µL of Fluoro-Development solution was added and incubated for 4 minutes before measuring fluorescence at 530Ex/590Em.

### Nuclear Protein Extraction and Quantification

A549 Transwell inserts were exposed to 0 and 750 µg mL^-1^ NiO or NiCl_2_ based on nickel content using a VITROCELL Cloud Alpha 6 exposure system. Following exposure, the co-culture was assembled and incubated for 24 hours in a humidified incubator at 37 °C. Nuclear proteins from both A549 cells and cardiomyocytes were extracted separately using the EpiQuik Nuclear Extraction Kit (EpigenTek, OP-0002) according to the manufacturer’s guidelines. Briefly, cells were trypsinized and centrifuged at 1000 rpm for 5 minutes. Pellets were resuspended in Pre-Extraction Buffer supplemented with dithiothreitol (DTT) and protease inhibitor cocktail (PIC) at a 1:1000 ratio. After a 10-minute incubation on ice, samples were vortexed and centrifuged at 12,000 rpm for 1 minute. The supernatant was discarded, and pellets were resuspended in Extraction Buffer containing DTT and PIC at the same ratio. Samples were incubated on ice for 15 minutes with vortexing every 3 minutes, followed by sonication (10 seconds, three cycles). Nuclear extracts were obtained by centrifugation at 14,000 rpm for 10 minutes, and the resulting supernatant was collected.

Protein concentration was determined using the Bio-Rad Protein Assay Dye Reagent Concentrate (Bio-Rad, 5000006) with BSA (Bio-Rad, 5000007) as the standard. In a 96-well plate, 5 µL of each sample or standard was mixed with 100 µL of 20 % dye reagent in ultrapure water. After 5 minutes of incubation, absorbance was measured at 595 nm, and protein concentrations were calculated from a BSA standard curve ranging from 15 to 1000 µg mL^-1^.

### DNA methyltransferase activity

The enzymatic activity of DNA methyltransferases (DNMTs) in A549 cells and hiPSC-CM was quantified using the EpiQuik DNMT Activity/Inhibition Assay Ultra Kit (EpigenTek, P-3010). Nuclear extracts (5 µL) were combined with 45 µL of working buffer in each well and incubated for 120 minutes at 37 °C. Wells were washed three times with the provided wash buffer, after which 50 µL of capture antibody were added and incubated for 60 minutes at room temperature. Following another three washes, 50 µL of detection antibody solution was added and incubated for 30 minutes. The wells were washed four times and incubated with 50 µL of enhancer solution for 30 minutes. After five final washes, 50 µL of Fluorescence Development Solution was added, and fluorescence was recorded after 3 minutes at 530Ex/590Em using a microplate reader.

### Comet assay

DNA strand breaks and oxidative DNA lesions, indicated by 8-oxoguanine, were assessed in A549 cells and hiPSC-CMs using the comet assay as previously described (Di Bucchianico et al., 2017). After 4 and 24 hours of exposure and co-culture assembly, A549 cells and hiPSC-CMs were harvested and frozen separately. For the comet assay, thawed cells were centrifuged at 200 ×g for 10 minutes and resuspended in cold DPBS. A 15-µL aliquot of each cell suspension was mixed with 70 µL of 1 % low-melting agarose (Fisher BioReagents, BP165). On ice-chilled slides pre-coated with 0.5 % agarose (Fisher BioReagents, BP164), 15-µL droplets of the mixtures were applied in duplicate onto two slides—one for the alkaline comet assay and the other for the FPG-modified comet assay. Slides were kept on ice until the agarose solidified, then incubated for 60 minutes at 4 °C in lysis buffer containing 2.5 M NaCl (Thermo Fisher Scientific, 207790050), 0.1 M EDTA (Fisher BioReagents, BP120), 10 mM Tris (Sigma-Aldrich, T4661), and 10 µg mL^-1^ Triton X-100 (Sigma, T8787). After lysis, slides were washed three times with reaction buffer (pH 8) containing 0.5 mM EDTA, 40 mM HEPES (Thermo Fisher Scientific, 172570250), 0.2 mg mL^-1^ BSA (Sigma-Aldrich, A2153), and 0.1 M KCl (Fisher BioReagents, BP366). For the FPG-modified comet assay, slides were incubated with FPG enzyme (New England Biolabs, M0240S) diluted 1:30,000, while corresponding alkaline comet slides were incubated with reaction buffer. All slides were incubated for 30 minutes at 37 °C, followed by cooling at 4 °C for 10 minutes. They were then immersed in electrophoresis buffer (pH 13) containing 1 mM EDTA and 300 mM NaOH (Fisher BioReagents, BP359) and for 40 minutes to allow DNA unwinding. Slides were transferred to an electrophoresis tank containing ice-cold electrophoresis buffer, and electrophoresis was performed at 20 V and 300 mA for 20 minutes. Afterwards, slides were washed twice with cold 0.5 M Tris and twice with cold ultrapure water before being air-dried for 2 days. For staining, 80 µL of SYBR Gold (Thermo Scientific, S11494) diluted 1:10,000 in 1× TE buffer (Santa Cruz Biotechnology, sc-296654) was applied to each slide and covered with a coverslip. Comets were imaged using a fluorescence microscope, and DNA damage was quantified using CometScore 2.0. Three independent biological replicates were analyzed, with 100 comets scored per replicate.

### Microelectrode Array Measurement

For the short-term measurement of the cardiac field potentials, the insert was first removed and then reinserted. Subsequent to a 15-minute equilibration period on the pre-heated headstage, the baseline signals were measured and the corresponding parameters were adjusted. The measurements were generally performed under ambient atmosphere at a temperature of 37 °C and a sampling rate of 10 kHz. Measurements were taken with a MEA2100 system and Cardio2D software (Multi-Channel Systems, Reutlingen, whereas the analysis was conducted with Cardio2D+ software (Multi-Channel Systems). The field potential (FP) frequency and field potential duration (FPD) was analysed on 5 out of 60 electrodes, whereas the conduction velocity was calculated on the area automatically.

### Statistical analysis

For medium selection and cell compatibility assessments, data were derived from two independent experiments and are presented as mean ± standard deviation (SD). All subsequent experiments involving nickel exposure were conducted in at least three independent biological replicates, each including two technical replicates. Data from these experiments are expressed as mean ± standard error of the mean (SEM). Graphical representation and quantitative analyses of cytotoxicity, metabolic activity, cytokine profiling, comet assay results, epigenetic endpoints and MEA analysis were performed using GraphPad Prism for Windows (GraphPad Software 11.0.1). Statistical analyses were conducted using one-way analysis of variance (ANOVA) followed by Bonferroni’s multiple-comparisons test.

## Results

### Optimization of Co-Culture Conditions and Biocompatibility between A549 Cells and hiPSC-CMs

Prior to nickel exposure experiments, it was essential to establish cultivation conditions that support both A549 cells and hiPSC-CMs without compromising their viability or functional characteristics. To identify an optimal co-culture medium, three formulations were evaluated: A549 medium, hiPSC-CM medium, and a 1:1 (v/v) mixture of both. Cytotoxicity in each cell type was assessed at 4, 24, and 48 hours after co-culture settings.

A549 cells exhibited comparable cytotoxicity levels when cultured with the different medium or their combination for 4 and 24 hours (Figure 1a). After 48 hours, the cytotoxicity increases in all conditions due to the continued proliferation and accumulation of dead cells without medium change.

**Figure 1.**
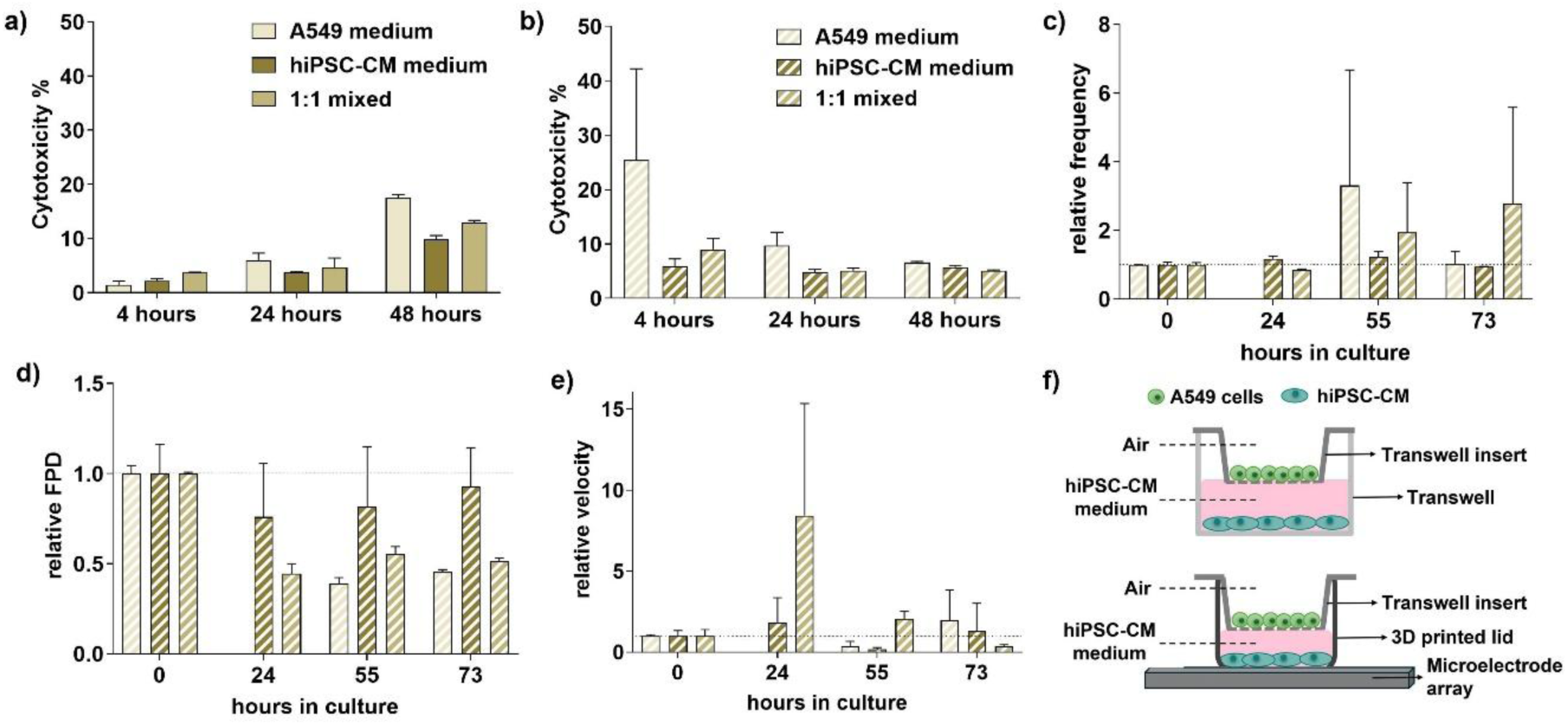
Validation of media compatibility. Cytotoxicity of A549 (a) and hiPSC-CM (b) cells cultured in A549 medium, hiPSC-CM medium, or a 1:1 mixture of both media for 4, 24, and 48 hours. hIPSC-CMs electrophysiological response to named media conditions with regard to beat frequency (c), FPD (D) and conduction velocity (e). Schematic representation of co-culture model of alveolar epithelial A549 cells and human induced pluripotent stem cell-derived cardiomyocytes (hiPSC-CMs) in transwell and on top of microelectrode array (f). Data are presented as mean ± SD from two independent experiments.

In contrast, hiPSC-CMs displayed a marked cytotoxic response to A549 medium as early as 4 hours post-exposure (Figure 1b). This indicates that this medium is unsuitable for cardiomyocyte culture, even though cytotoxicity was reduced to control levels at 24- and 48-hours incubation time. Functional electrophysiological measurements further supported this conclusion. hiPSC-CMs cultured in A549 medium or the mixed medium showed an elevated field potential frequency and significantly shortened field potential duration (FPD) especially after 55 and 73 hours (Figures 1c and 1d), suggesting medium-induced stress or altered cardiac electrophysiology. The signal propagation (velocity) is most stable under CM medium (Figure 1e), while the 1:1 mixed medium appears to temporarily increase the conduction velocity. Collectively, these findings indicate that CM medium is the most compatible and physiologically stable option for maintaining both cell types in a co-culture system (Figure 1f).

To ensure that the two cell types could be co-cultured without negatively influencing each other, biocompatibility was evaluated by measuring cytotoxicity, metabolic activity, and cardiomyocyte field potential characteristics in mono-cultures and co-cultures. Co-culturing for 24 hours did not induce significant cytotoxicity in either A549 cells or hiPSC-CMs, with values remaining below 2% cytotoxicity (Figure 2a). Metabolic activity was equally unaffected (Figure 2b), indicating preserved cellular function. Additionally, the field potential frequency of hiPSC-CMs remained stable at 0, 4, and 24 hours after co-culture formation (Figure 2c) as well as the field potential duration (Figure 2d). However, the field potential amplitude was reduced in co-cultures (Figure 2e), which may indicate an influence of the insert/insert removal. Furthermore, the conduction velocity (Figure 2f) of the CM also demonstrates differences between the mono- and co-cultures over time. While the conduction velocity of the monoculture increases, indicative of enhanced synchronisation of the cell layer, it exhibits a temporary decrease in the co-culture following a 4-hour period, subsequently returning to its baseline state. As a consequence, sufficient distance between adherent hiPSC-CM cells and inserts must always be ensured in subsequent experiments.

**Figure 2.**
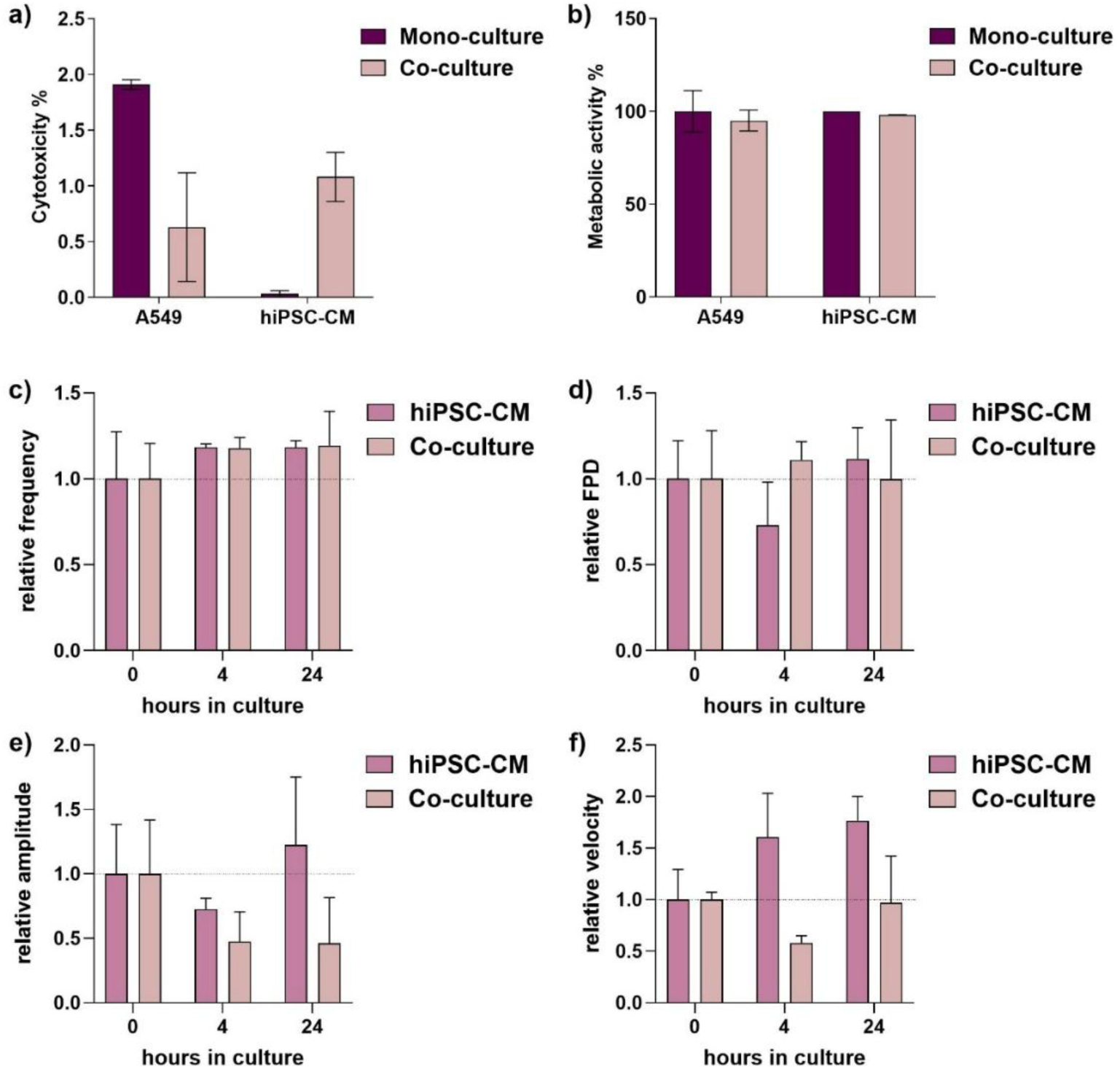
Validation of co-culture cultivation conditions. Cell type compatibility in the co-culture model after 24 hours was assessed by cytotoxicity (a) and metabolic activity (b) on A549 and hiPSC-CM. Cardiac field potential analysis on frequency (c), FPD (d), amplitude (e) and conduction velocity (f) were performed on CM monocultures and co-cultures with A549 inserts before, 4 and 24 hours after starting the co-culture. Data are presented as mean ± SD from two independent experiments. MEA data are normalized to the respective baseline values.

These results demonstrate that A549 cells and hiPSC-CMs are biocompatible and can be co-cultured in CM medium without disturbing baseline viability or cardiomyocyte function.

### Cytotoxic Responses of Co-cultures to Particulate NiO and Ionic NiCl_2_ Nickel

The co-culture model was exposed to 3 concentrations of NiO nanoparticles or NiCl_2_ at ALI and then incubated for 4 or 24 hours. In the following, the basolateral medium shared by A549 and hiPSC-CM cells was analysed. Both NiO and NiCl_2_ induced slight cytotoxicity after 4 hours of exposure (Figure 3a). After 24 hours, exposure to 280 and 840 ng Ni cm^-2^ of NiO led to a significant increase in cytotoxicity (Figure 3b). No clear dose-dependent pattern emerged at either timepoint, which may reflect concentration-dependent changes in particle agglomeration, or activation of antagonistic cellular responses.

**Figure 3.**
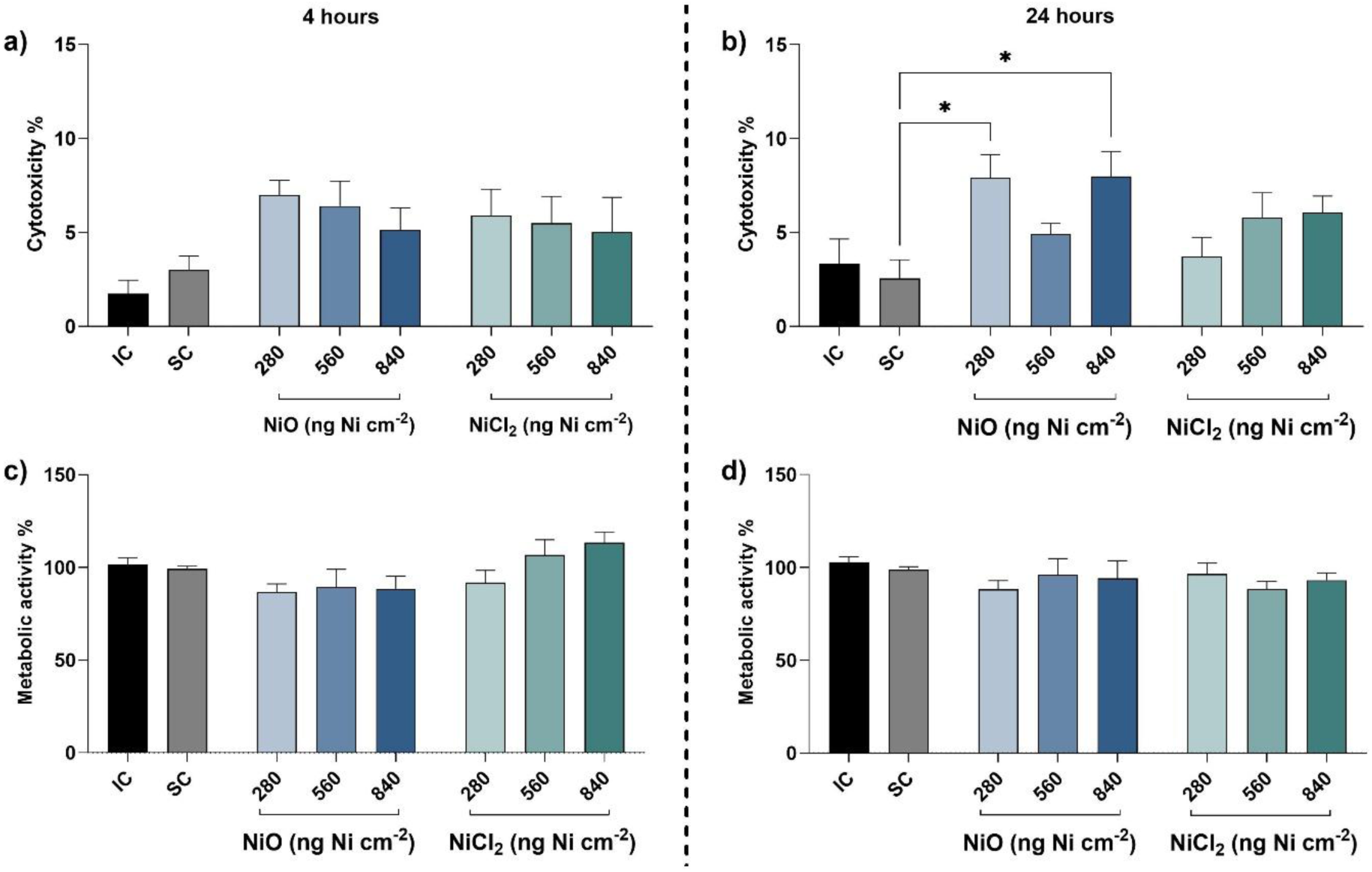
Cytotoxicity and metabolic activity in A549/hiPSC-CM co-cultures following NiO and NiCl_2_ ALI exposure. Cytotoxicity measured at 4 hours (a) and 24 hours (b) following exposure. Metabolic activity assessed at 4 hours (c) and 24 hours (d) following exposure. *p < 0.05, **p < 0.01.

Metabolic activity assessed under the same exposure conditions showed no significant alterations at 4 or 24 hours (Figures 3c and 3d). Collectively, these results indicate that although nickel exposure, particularly to NiO, can induce cytotoxic responses, overall cellular metabolic activity remains largely unaffected. Furthermore, the magnitude of toxicity did not differ markedly between particulate and ionic nickel. Furthermore, the magnitude of toxicity did not differ markedly between particulate and ionic nickel.

### Nickel-Induced Inflammatory Activation Assessed by Cytokine Release

Cytokine release into the shared medium of A549 cells and hiPSC-CMs was analyzed to evaluate whether secondary, cytokine-mediated effects on cardiomyocytes could occur following nickel exposure. To this end, five pro-inflammatory cytokines (IL-1β, IL-6, IL-8, IL- 18, TNF-α) and one anti-inflammatory cytokine (IL-10) were quantified 4- (Figure 4) and 24-hours (Figure 5) post exposure.

**Figure 4.**
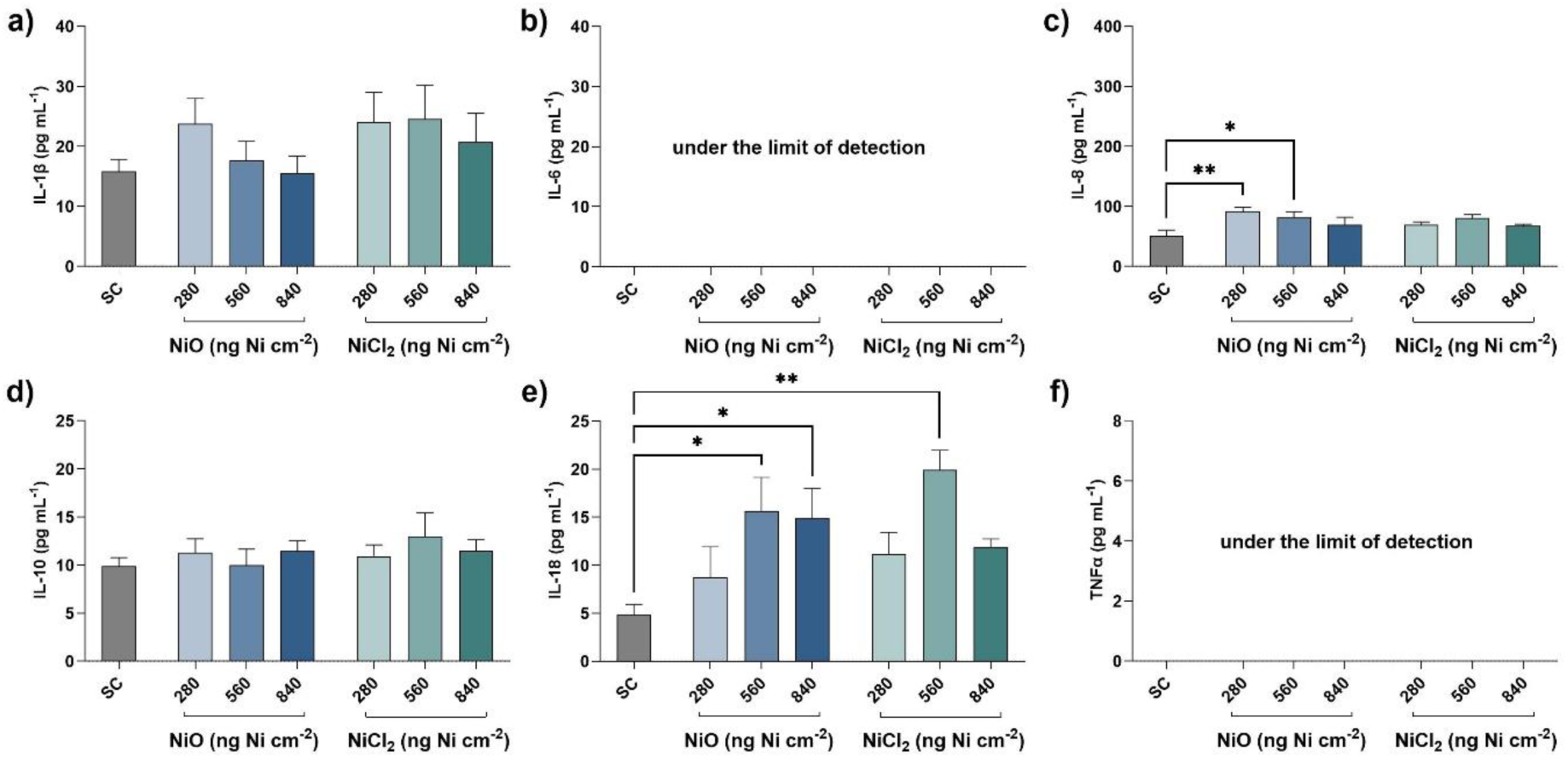
Cytokines levels of the co-culture 4 hours following NiO and NiCl_2_ ALI exposure. IL-1β (a), IL-6 (b), IL-8 (c), IL-10 (d), IL-18 (e), and TNF-α (f). *p < 0.05, **p < 0.01.

**Figure 5.**
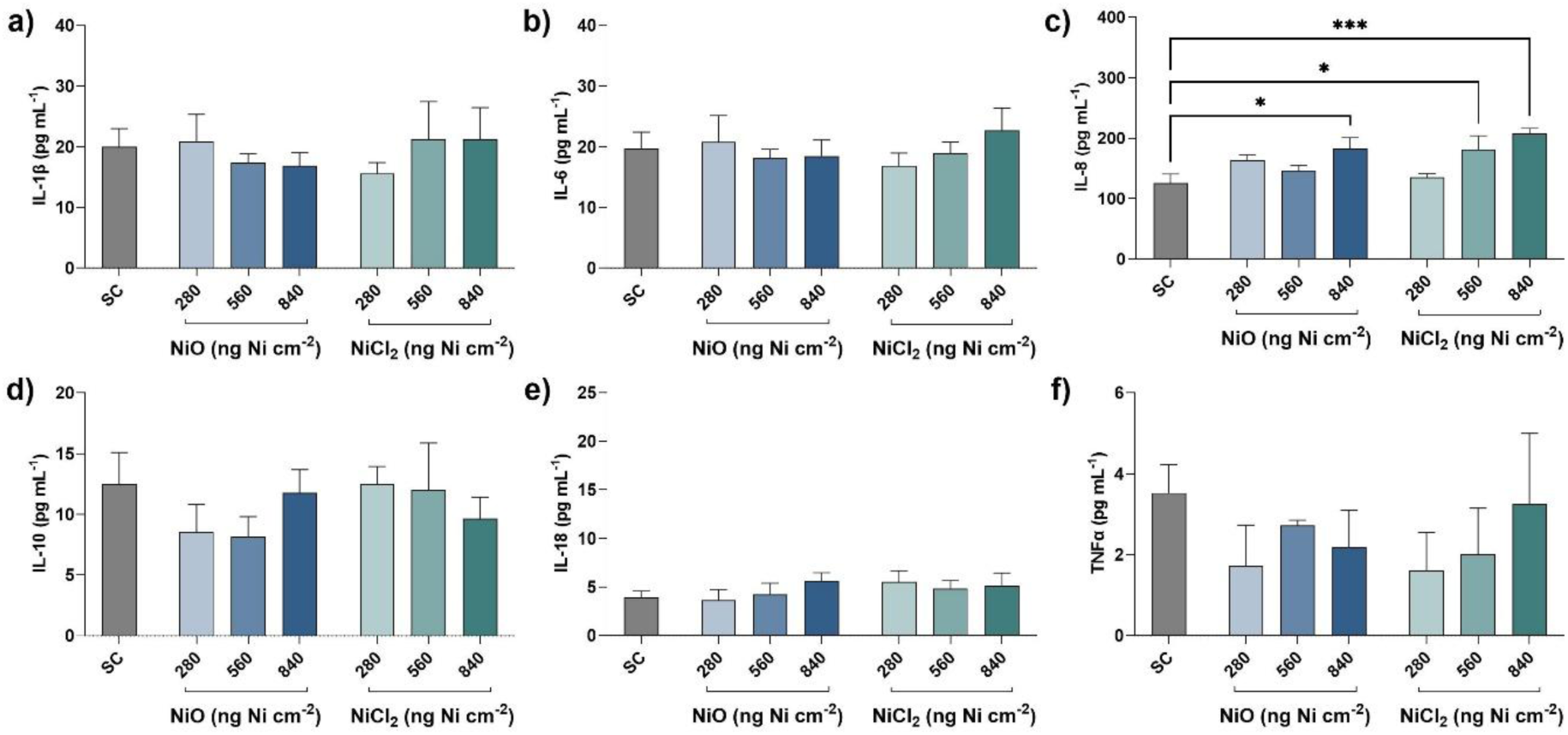
Cytokines levels of the co-culture 24 hours following NiO and NiCl_2_ ALI exposure. IL-1β (a), IL-6 (b), IL-8 (c), IL-10 (d), IL-18 (e), and TNF-α (f). *p < 0.05, ***p < 0.001.

Following NiO exposure, IL-8 secretion increased after 4 hours compared with the solvent control (Figure 4c). In addition, IL-18 levels were elevated following exposure to both NiO and NiCl₂ (Figure 4e), indicating an early inflammatory response to both nickel forms. Across all exposure conditions, IL-8 levels increased substantially between 4 and 24 hours, consistent with time-dependent accumulation of this cytokine (Figure 5c). However, no marked differences were observed between NiO and NiCl₂, suggesting that IL-8 induction was not strongly dependent on the initial nickel form under the applied exposure conditions. In contrast, the early increase in IL-18 observed at 4 hours was no longer evident at 24 hours (Figure 5e), indicating a transient activation of this inflammatory response.

To determine whether the co-culture system itself contributed to IL-8 production, cytokine levels were assessed in A549 monocultures, hiPSC-CM monocultures, and their co-culture. The cumulative IL-8 levels of the monocultures closely matched those observed in the co-culture, indicating that co-cultivation does not intrinsically trigger IL-8 signalling. As expected, A549 cells were the primary source of IL-8, contributing approximately 72–73% of the total cytokine levels at both time points.

No significant changes were detected for IL-1β, IL-6, IL-10, or TNF-α at either time point following exposure to NiO or NiCl₂. This suggests that, under the applied conditions, nickel exposure selectively induces IL-8 and IL-18 without broadly activating additional pro- or anti-inflammatory cytokine responses.

### Basolateral Translocation of Nickel Across the ALI Co-Culture Membrane Following NiO and NiCl_2_ Exposure

The established co-culture model was exposed to three concentrations of NiO nanoparticles or NiCl_2_ at ALI for 4 or 24 hours before the collection of basolateral medium for nickel quantification. Nickel translocation across the Transwell membrane was quantified by atomic absorption spectroscopy to estimate the extent of basolateral exposure of hiPSC-CMs and thereby the potential for direct cellular contact with nickel. This is relevant for distinguishing indirect paracrine effects from possible direct exposure of the cardiac compartment following ALI treatment and nickel translocation.

As shown in Table 1, nickel was detectable in the basolateral medium after exposure to both NiO nanoparticles and NiCl_2_ at 4 and 24 hours. The data did not follow a strict dose-response relationship for NiO nanoparticles. Following NiO exposure, basolateral nickel levels remained relatively stable between 4 and 24 hours, suggesting a balanced bioavailability of nickel from the particulate form with no-time dependent increase in translocation. In contrast, NiCl_2_ showed a clear increase in basolateral nickel concentration over time, indicating progressive diffusion of ionic nickel across the membrane and greater bioavailability to the underlying cardiomyocytes at later time points. Calculation of translocation efficiency further revealed that only a small fraction of the deposited nickel reached the basolateral compartment, ranging from approximately 1–8% of the deposited dose. Notably, the percentage of translocated nickel was not directly proportional to the deposited nickel dose for either NiO or NiCl₂.

**Table 1:** Nickel translocation to the basal medium 4- and 24-hours post-exposure.

|  |  |  |  |  |
| --- | --- | --- | --- | --- |
|  | **4 hours** |  | **24 hours** |  |
|  | **Basolateral Ni concentration ($\mu\text{g L}^{-1}$)\*** | **Percentage of translocated Ni** | **Basolateral Ni concentration ($\text{ng mL}^{-1}$)\*** | **Percentage of translocated Ni** |
| **IC** | 2.07 ± 0.03 | NA | 1.85 ± 0.07 | NA |
| **SC** | 4.56 ± 1.42 | NA | 3.65 ± 0.80 | NA |
| **NiO ($\text{ng cm}^{-2}$)** |  |  |  |  |
| **280** | 15.84 ± 7.81 | 7.76 ± 3.83 | 12.57 ± 1.42 | 6.16 ± 0.70 |
| **560** | 8.26 ± 2.41 | 2.03 ± 0.59 | 11.28 ± 2.86 | 2.77 ± 0.70 |
| **840** | 17.73 ± 8.43 | 2.90 ± 1.38 | 6.23 ± 1.02 | 1.02 ± 0.17 |
| **NiCl<sub>2</sub> ($\text{ng cm}^{-2}$)** |  |  |  |  |
| **280** | 6.99 ± 1.31 | 3.43 ± 0.64 | 9.63 ± 2.35 | 4.72 ± 1.15 |
| **560** | 13.96 ± 4.90 | 3.42 ± 1.20 | 35.21 ± 12.82 | 8.63 ± 3.14 |
| **840** | 17.71 ± 6.54 | 2.89 ± 1.07 | 45.00 ± 32.32 | 7.35 ± 5.28 |
| Data are presented as mean ± SEM. | | | | |
| * Nickel concentrations in the basolateral compartment were quantified by atomic absorption spectroscopy (AAS). |  |  |  |  |
| ** Total basolateral nickel mass (ng) was calculated based on the measured nickel concentration and the basolateral medium volume (1.5 mL). |  |  |  |  |

Together, these findings demonstrate distinct translocation behaviours for particulate and ionic nickel. NiO appears to function as a relatively stable reservoir with limited release, whereas NiCl_2_ provides a more rapidly and continuously increasing basolateral nickel exposure. This difference is important for interpretation of downstream biological effects, suggesting that ionic nickel is more likely to contribute more to direct cardiomyocyte exposure after 24 hours, whereas nanoparticulate nickel may exert its effects through slower release dynamics and indirect intercellular signalling pathways.

### DNA Strand Breaks and Oxidative DNA Damage Following NiO and NiCl_2_ Exposure

DNA damage in A549 cells and hiPSC-CMs was evaluated separately using the comet assay, with DNA strand breaks quantified as the percentage of DNA in the tail. Baseline levels of DNA damage were lower in A549 cells than in hiPSC-CMs (Figure 6a–d), suggesting a higher intrinsic susceptibility of hiPSC-CMs to DNA strand breaks.

**Figure 6.**
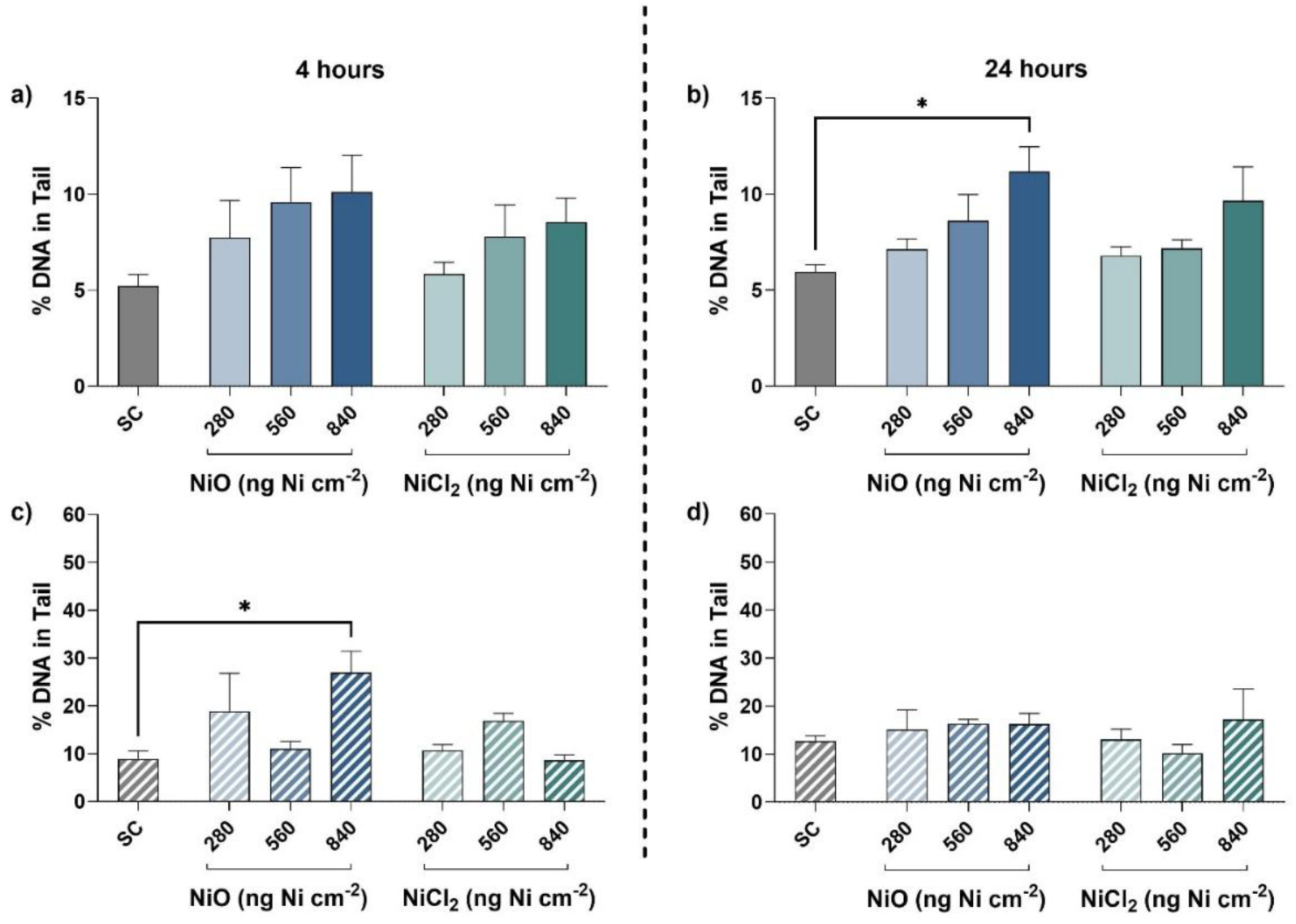
DNA strand breaks assessed by Comet assay in A549/hiPSC-CM co-cultures following NiO and NiCl_2_ ALI exposure. Percentage of DNA in tail, indicating DNA strand breaks, in A549 cells at 4 hours (a) and 24 hours (b), and in hiPSC-CM at 4 hours (c) and 24 hours (d). *p< 0.05.

In A549 cells, no significant increase in DNA strand breaks was observed after 4 hours of exposure (Figure 6a). However, after 24 hours, exposure to 840 ng Ni/cm² of NiO resulted in a significant elevation in DNA strand breaks (Figure 6b). In contrast, hiPSC-CMs exhibited a distinct temporal pattern: marked DNA damage was detected at 4 hours following exposure to 840 ng Ni/cm² of NiO (Figure 6c), whereas this effect was no longer evident at 24 hours (Figure 6d). These findings indicate cell type-specific differences in sensitivity to nickel-induced genotoxicity, as well as distinct temporal patterns of damage induction and potential repair responses.

Analysis of oxidative DNA lesions revealed a slight increase in oxidative damage in A549 cells at 4 hours across treatments (Figure 7a). This effect persisted at 24 hours for the two highest concentrations, while diminishing at the lowest concentration (Figure 7b). In hiPSC-CMs, no oxidative DNA damage was observed following NiO exposure at 4 hours; however, NiCl_2_ induced a significant increase at the highest concentration (Figure 7c). This pattern remained consistent at 24 hours, with NiO showing no effect and NiCl_2_ continuing to elevate oxidative DNA damage (Figure 7d).

**Figure 7.**
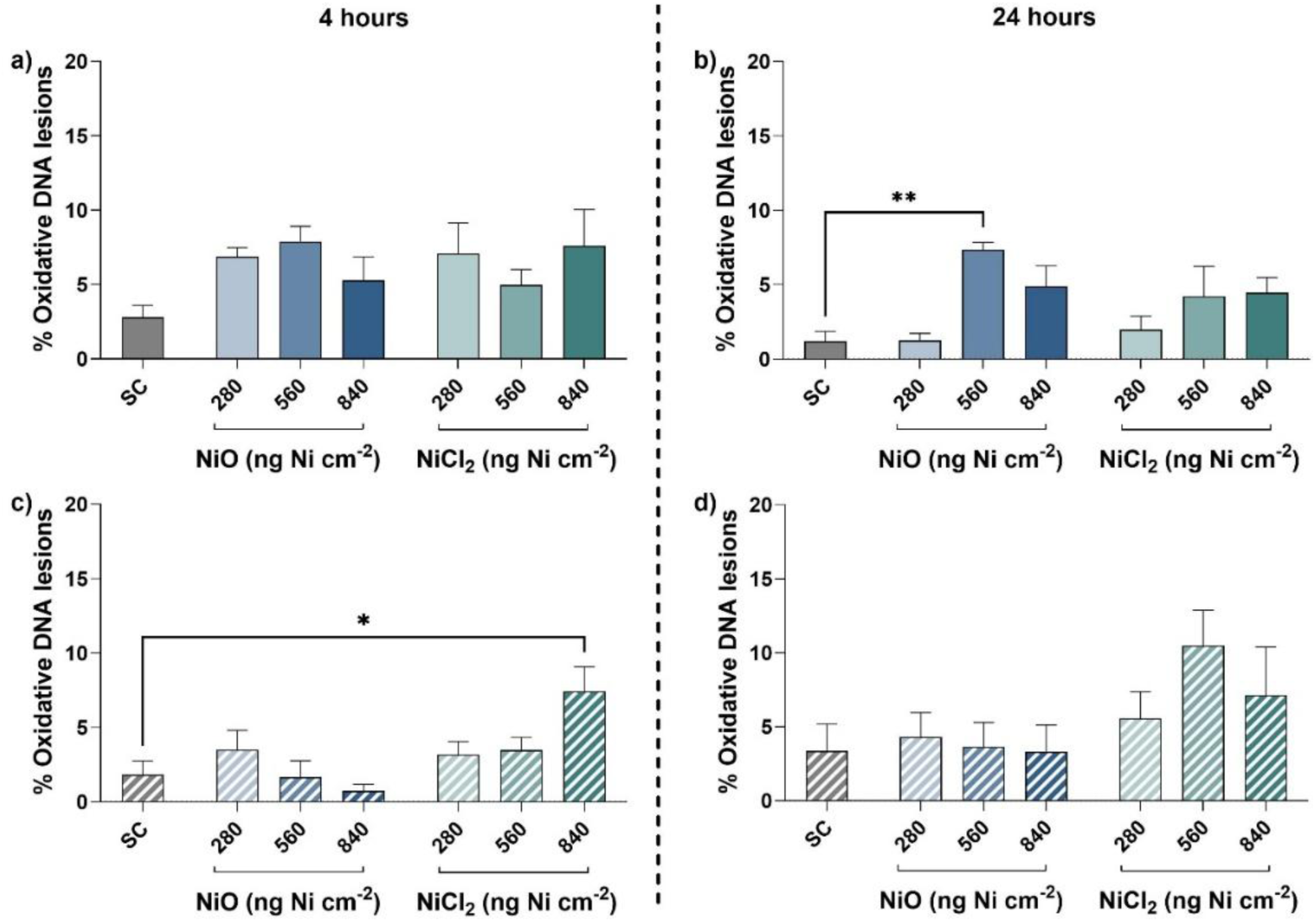
Oxidative DNA lesions assessed by FPG-modified Comet assay in A549/hiPSC-CM co-cultures following NiO and NiCl_2_ ALI exposure. Oxidative DNA lesions percentage in A549 cells at 4 hours (a) and 24 hours (b), and in hiPSC-CM at 4 hours (c) and 24 hours (d). * p< 0.05, ** p< 0.01.

Neither nickel compound produced a uniform dose-response relationship across all endpoints, except for DNA strand breaks in A549 cells. Notably, compound-specific differences emerged: NiO more prominently induced DNA strand breaks in both cell types, whereas NiCl_2_ particularily elicited oxidative DNA damage in hiPSC-CMs. In A549 cells, NiCl_2_ did not result in significant DNA damage under the tested conditions.

### Global DNA Methylation Alterations and DNMT Activity in Response to Nickel Exposure

Epigenetic modifications post exposure were investigated by global cytosine methylation levels and the activity of DNA methyltransferases (DNMTs) in both A549 cells and hiPSC-CMs separately. The global cytosine methylation levels were assessed to evaluate whether Nickel compounds can cause an epigenetic shift. The concentration 840 ng Ni cm^-2^ was used for exposure with both NiO and NiCl_2_, as this concentration caused a predictable change in all the previously reported data. Neither treatment significantly reduced global 5-methylcytosine levels in A549 cells (Figure 8a). In contrast, hiPSC-CMs exhibited a significant reduction, with levels decreased by 3.8-fold and 3.6-fold following exposure to NiO and NiCl_2_, respectively. Additionally, hiPSC-CMs displayed higher baseline methylation levels compared with A549 cells, which may be attributable to the cancerous nature of A549 cells and the non-proliferating and differentiated status of hiPSC-CMs.

**Figure 8.**
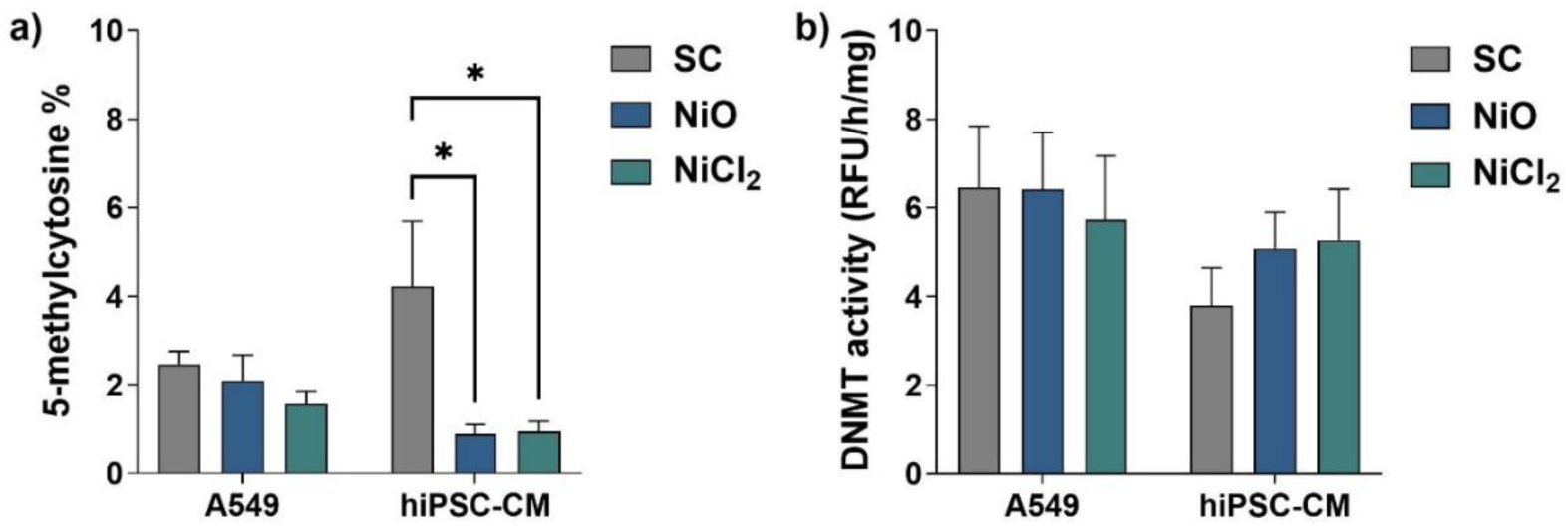
Epigenetic modifications in A549/hiPSC-CM co-cultures following 24 hours ALI exposure to 840 ng Ni cm^-2^ of NiO or NiCl_2_. Levels of 5-methylcytosine in A549 cells and hiPSC-CMs (a). DNA methyltransferase (DNMT) activity in A549 cells and hiPSC-CMs (b). *p < 0.05.

We investigated the activity of DNMTs in A549 cells and hiPSC-CMs upon exposure to 840 ng Ni cm^-2^ of both nickel species. The activity of DNMTs in both cell types was not significantly altered (Figure 8b), which indicates that the decrease of methylation levels in cells is not caused by a decrease in the activity of DNMTs. Still, both forms of Nickel were not different from each other in terms of global methylation and DNMT activity.

### Functional Cardiac Effects Following Exposure to Nickel Compounds

Nickel-exposed A549 cells were co-cultured with hiPSC-CMs for up to 24 hours, with functional measurements obtained at 0, 4, and 24 hours after co-culture assembly. Cardiac electrophysiological activity was assessed using microelectrode arrays, including field potential frequency, FPD, and conduction velocity across the cardiomyocyte network. For both NiO and NiCl_2_, field potential frequency remained largely stable from baseline (0 hours) to 24 hours, with only a slight reduction observed following NiO exposure at 4 and 24 hours (Figure 9a–c). Similarly, FPD values showed no substantial deviations from the solvent control across most conditions, indicating minimal effects on repolarization dynamics (Figure 9d–f). In contrast, analysis of conduction velocity revealed transient alterations. An increase in conduction velocity was observed immediately after co-culture assembly (0 hours) following NiCl_2_ exposure and at 4 hours following NiO exposure (Figure 9g–h). However, by 24 hours, conduction velocity decreased to levels comparable to the solvent control, indicating that these effects were not sustained over time (Figure 9i). This suggests a short-lived modulation of electrical signal propagation across the cardiomyocyte layer.

**Figure 9.**
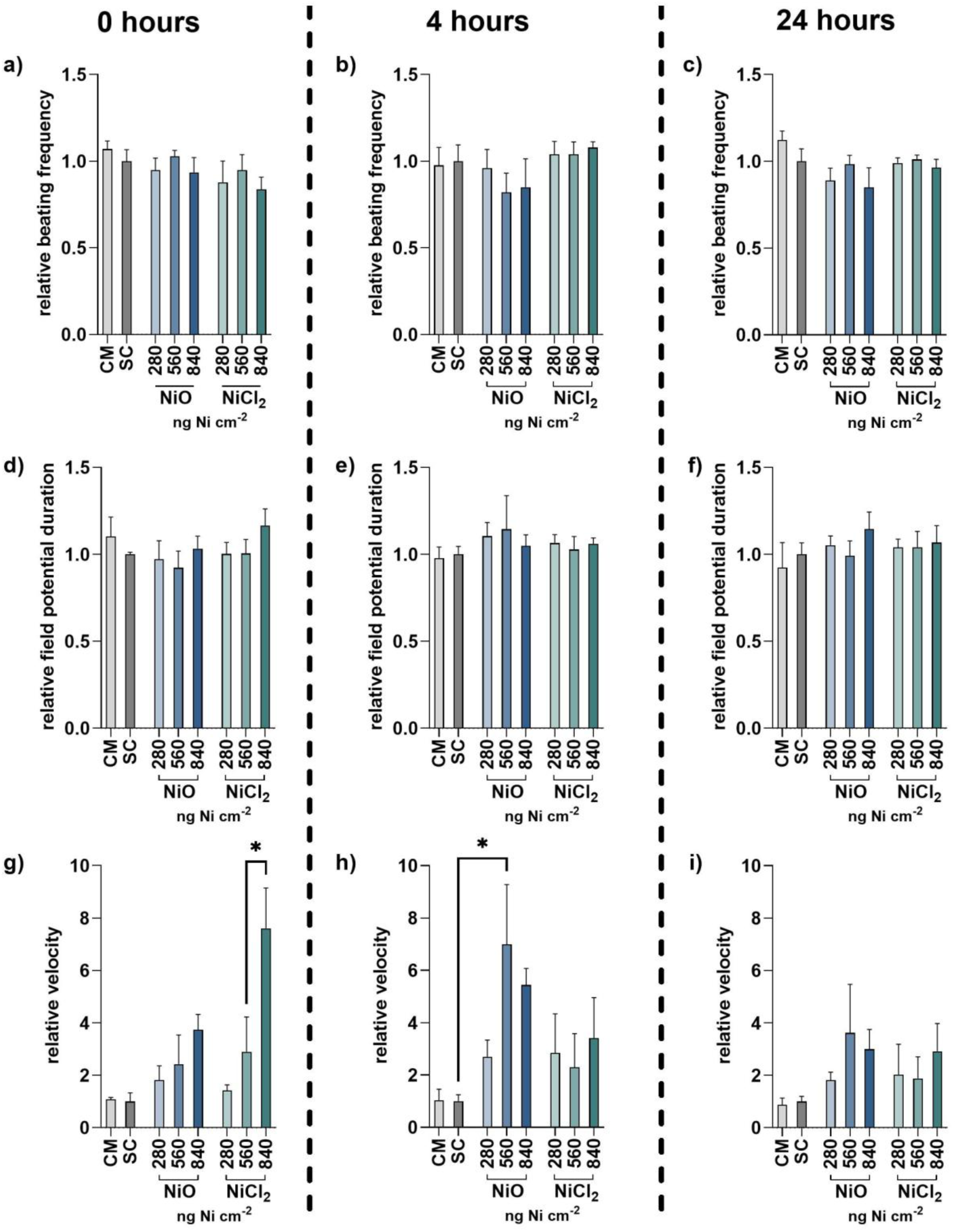
Effects of nickel compound exposure on electrophysiological properties of hiPSC-CMs following co-culture with A549 cells. Relative beating frequency of hiPSC-CMs at 0 hours (a), 4 hours (b), and 24 hours (c). Relative field potential duration (FPD) at 0 hours (d), 4 hours (e), and 24 hours (f). Relative conduction velocity at 0 hours (g), 4 hours (h), and 24 hours (i). Data are normalized to the respective solvent control (SC) at each time point. CM control represents hiPSC-CM monoculture. *p < 0.05.

## Discussion

This study establishes a human-relevant ALI lung-heart co-culture platform that enables the investigation of both primary pulmonary effects and secondary cardiac responses following inhalation-relevant exposure to nickel in particulate (NiO) and ionic (NiCl_2_) forms. By integrating epithelial exposure at the ALI with functional cardiomyocyte readouts, the model addresses a critical gap between existing in vitro systems and complex in vivo studies. The results of this study demonstrate that nickel exposure induces cytotoxicity and inflammatory response characterized by the release of IL-8 and IL-18 in the co-culture system. In A549 cells, NiO exposure resulted in a delayed increase in DNA strand breaks and oxidative DNA lesions. In contrast, hiPSC-CMs exhibited an early but transient increase in DNA strand breaks and oxidative DNA damage following exposure to NiO and NiCl_2_, respectively. Furthermore, both nickel forms induced a reduction in global DNA methylation levels in hiPSC-CMs after 24 hours. Electrophysiological analysis revealed transient changes in cardiomyocyte function, with increased relative conduction velocity observed following ionic nickel exposure at 4 hours and NiO exposure at 24 hours, while other functional parameters remained largely unaffected.

A key prerequisite for interpreting these findings was the validation of compatible co-culture conditions. Cardiomyocyte medium was essential to maintain stable electrophysiological behaviour in hiPSC-CMs. This selectivity aligns with the known metabolic demands of cardiomyocytes, which require insulin support for proper contractile function (Yu et al., 2006). Co-culture itself did not induce significant cytotoxicity or metabolic impairment, confirming that the observed effects are exposure-driven rather than model-induced effects. Minor alterations in electrophysiological parameters could be controlled by running a parallel hiPSC-CM monoculture as a reference throughout all MEA measurements.

Evaluating the co-culture model using a well-characterized toxicant represents a critical step in validating its functionality and biological responsiveness. Nickel was therefore selected in both its nanoparticulate (NiO) and ionic (NiCl_2_) forms to assess the system’s capacity to capture relevant toxicological outcomes. As expected, nickel exposure induced cytotoxicity in the co-culture, consistent with previous reports demonstrating its adverse effects in both pulmonary and cardiac cell models (Forti et al., 2011; Lou et al., 2013; Mo et al., 2024). Despite their distinct forms, NiO and NiCl_2_ elicited comparable levels of cytotoxicity. This apparent similarity likely reflects distinct kinetic profiles, whereby NiO functions as a slowly dissolving reservoir of nickel inside the cell while also exerting particle-related biological effects (Latvala et al., 2016; 2018; Gliga et al., 2020; Zhou et al., 2025), whereas NiCl_2_ delivers an immediately bioavailable ionic form. However, the reported intracellular dissolution of NiO nanoparticles and their rapid agglomeration (Latvala et al., 2016; Di Bucchianico et al., 2018; Shinohara et al., 2017), could drive downstream effects, and complicate direct comparisons between particulate and ionic forms.

Both NiO and NiCl_2_ have been reported to induce inflammatory responses (Capasso et al., 2014; Du et al., 2025); therefore, we assessed the release of six cytokines in the co-culture following nickel exposure. A key finding of this study is the selective induction of IL-8 and IL-18, whereas no significant changes were observed for IL-1β, IL-6, IL-10, or TNF-α. IL-8 is a central chemokine involved in neutrophil recruitment and is widely recognized as a sensitive marker of epithelial stress in response to metal exposure (Hammond et al., 1995; Honda et al., 2015). In contrast, IL-18 is a pro-inflammatory cytokine produced by cardiomyocytes and has been associated with myocardial injury and adverse cardiac remodelling processes (Wang et al., 2008; Suetomi et al., 2018). The absence of a broader cytokine response suggests that the applied exposure conditions elicit a targeted, rather than generalized, inflammatory profile. Mechanistically, this selective induction may reflect preferential activation of NF-κB-dependent signalling pathways, as nickel exposure has previously been shown to stimulate IL-8 production through NF-κB activation (Freitas et al., 2010). Similarly, NF-κB signalling has been implicated in the regulation of IL-18 expression and inflammatory responses in cardiomyocytes (Chandrasekar et al., 2003; Wang et al., 2008).

Genotoxicity analysis revealed distinct cell type- and time-dependent responses, highlighting differences in susceptibility between pulmonary epithelial cells and differentiated cardiomyocytes. A549 cells exhibited delayed induction of DNA strand breaks, whereas hiPSC-CMs displayed an early but transient increase in damage, suggesting either a greater initial sensitivity followed by more efficient repair processes. The higher baseline levels of DNA strand breaks detected in hiPSC-CMs compared with A549 cells may additionally reflect the accumulation of ROS and DNA damage associated with the cellular stress of the differentiation process (Miller et al., 2020). In hiPSC-CMs, the more pronounced oxidative DNA damage induced by NiCl_2_ supports a role for soluble nickel ions in ROS-mediated genotoxicity. Soluble nickel species have been reported to promote ROS generation and DNA strand breaks (Stinson et al., 1992; Chen et al., 2010), consistent with the detected increase in 8-oxyguanine in hiPSC-CMs with ionic Ni exposure. In contrast, NiO appeared to contribute more prominently to DNA strand breaks, potentially reflecting intracellular retention of particles and the gradual release of nickel ions following endocytic uptake. These observations align with previous studies demonstrating that poorly soluble nickel compounds can exhibit enhanced carcinogenic effect (Pietruska et al., 2011) and DNA damage (Åkerlund et al., 2018; Di Bucchianico et al., 2018), potentially due to prolonged intracellular persistence.

Nickel has been shown to interfere with epigenetic regulation through multiple mechanisms, including alterations in DNA methylation (Zheng et al., 2022, 2023), non-coding RNAs expression (Wu et al., 2017), and histone modifications (Jose et al., 2014, 2018, 2019). In the present study, epigenetic analysis revealed a significant reduction in global DNA methylation in hiPSC-CMs, whereas A549 cells remained largely unaffected following nickel exposure. Notably, these changes occurred in the absence of altered DNA methyltransferase activity, suggesting the involvement of alternative epigenetic regulatory mechanisms, potentially including modulation of demethylation process by Ten-Eleven Translocation (TET) enzyme activity. Baseline global methylation levels were higher in hiPSC-CMs than in A549 cells, which may reflect the cancerous nature of A549 cells that commonly exhibit global hypomethylation patterns (Tajbakhsh et al., 2022), as well as the differentiated and largely non-proliferative phenotype of cardiomyocytes. The greater sensitivity of hiPSC-CMs to epigenetic perturbation may additionally be linked to their developmental and differentiated state, since cardiac lineage commitment and maintenance require tightly regulated DNA methylation patterns controlling suppression of non-cardiac genes and permanent cell-cycle arrest (Kim et al., 2021).

A major contribution of this study lies in linking pulmonary exposure to functional cardiac outcomes. Functional electrophysiological analysis demonstrated that indirect exposure to nickel induced transient alterations in cardiomyocyte activity, specifically an increase in conduction velocity at 0 hours following NiCl_2_ exposure and at 4 hours following NiO exposure, while beating frequency and field potential duration remained largely stable. This implies subtle changes in electrical propagation rather than overt electrophysiological dysfunction. By 24 hours, conduction velocity decreased to levels comparable to the solvent control, indicating that these effects were not sustained over time. These alterations could suggest disrupted ion channel kinetics (Izumi-Nakaseko et al., 2018). The lack of sustained effects at 24 hours suggests that either adaptive cellular responses were engaged or that the concentration of nickel or inflammatory mediators reaching the cardiomyocytes was insufficient to produce lasting dysfunction under the present experimental conditions.

Although nickel was consistently detected in the basolateral compartment, only a relatively small fraction of the deposited dose crossed the epithelial barrier. This suggests that most deposited nickel remained within the apical compartment, likely due to cellular uptake, intracellular retention, or association with particulate agglomerates. Nevertheless, the presence of nickel in the basolateral medium confirms translocation across the epithelial barrier under the applied ALI conditions and demonstrates the potential for direct cardiomyocyte exposure within the co-culture system. Notably, the translocation kinetics differed between the two nickel forms. NiO exhibited relatively stable basolateral concentrations at 4 and 24 hours, consistent with its behaviour as a poorly soluble particulate reservoir that gradually releases nickel ions over time (Schaumlöffel, 2012). In contrast, NiCl_2_ showed a time- and concentration-dependent increase in basolateral nickel levels, indicating continuous diffusion and increasing bioavailability of soluble nickel species. These findings suggest that, although both forms contribute to cardiomyocyte exposure, NiCl_2_ is likely to induce more direct cellular effects through rapid ion availability, whereas NiO-mediated responses may arise from a combination of particle-specific effects and gradual ion release. Furthermore, the biological responses observed in hiPSC-CMs are unlikely to be driven exclusively by direct nickel exposure, as indirect mechanisms involving inflammatory mediators, such as IL-8, and oxidative stress-related signalling originating from the pulmonary compartment may also contribute to the observed effects.

The absence of a consistent dose-response relationship across several endpoints observed in the present study likely reflects the complex behaviour of nickel under ALI exposure conditions and within the co-culture system. Biological responses are influenced not only by the nominal deposited dose, but also by factors such as particle agglomeration, dissolution kinetics, and cellular uptake efficiency, especially under ALI exposures (Lacroix et al., 2018). Particularly for NiO, the gradual dissolution of nanoparticles may result in delayed or fluctuating intracellular ion concentrations that do not scale linearly with the applied concentration. In addition, low-dose exposures may already be sufficient to activate stress-responsive signalling pathways, resulting in plateau-like responses at higher concentrations (Lenz et al., 2013). The co-culture configuration further adds biological complexity, as indirect signalling between pulmonary epithelial cells and cardiomyocytes may amplify or modulate responses independently of the initial exposure concentration. Variability in membrane translocation kinetics and cellular defense mechanisms, including antioxidant responses and DNA repair capacity, may also contribute to the non-linear patterns observed across endpoints.

Collectively, the data support a model in which nickel exposure initiates pulmonary inflammation, oxidative stress, and DNA damage, which propagate to cardiomyocytes via soluble mediators such as IL-8 and potentially oxidative stress mediators, while limited but measurable translocation of nickel across the epithelial barrier contributes to direct cardiac exposure. The ALI co-culture system presented here thus provides a powerful platform for dissecting inter-organ communication and reveals mechanistic links between inhaled metal exposure and cardiac dysfunction. This integrated approach overcomes key limitations of traditional animal models, while enabling functional assessment of cardiomyocyte electrophysiology and contractility. Future studies incorporating more complex lung models, primary epithelial cells, and vascular endothelium, will further refine the physiological relevance of this approach and improve mechanistic resolution of lung-heart toxicology. Additionally, direct quantification of nickel speciation in basolateral medium and inclusion of longer exposure durations would help distinguish persistent effects and clarify the relative contributions of particle-specific versus ion-mediated effects.

## Author contributions

Wesam Darwish: Methodology, investigation (lung cell experiments), formal analysis, visualization, and writing – original draft. Sophie Kussauer: Methodology, investigation (cardiomyocyte experiments), formal analysis, visualization, and writing – original draft. Mohammad Almasaleekh: Methodology, investigation (lung cell experiments), and writing – review & editing. Sebastiano Di Bucchianico, Ralf Zimmermann, and Robert David: Conceptualization, methodology, validation, formal analysis, supervision, project administration, funding acquisition, and writing – review & editing.

## Acknowledgments

The authors would like to express their sincere and heartfelt gratitude to Sabine Haack and Andrea Schaarschmidt (University of Rostock) for their outstanding contributions to this work. Their expertise and dedication in performing the atomic absorption spectrometry measurements were essential for generating the nickel translocation data. In addition, their valuable input in developing, optimizing, and describing the methodological approach greatly strengthened the quality of this study.

## Data Availability

All data generated or analyzed during this study are available from the corresponding author upon reasonable request.

## Notes

### Competing Interest Statement

The authors have declared no competing interest.

## References

1. Åkerlund E, Cappellini F, Di Bucchianico S, et al. (2018) ‘Genotoxic and mutagenic properties of Ni and NiO nanoparticles investigated by comet assay, γ-H2AX staining, Hprt mutation assay and ToxTracker reporter cell lines.’, Environ Mol Mutagen, 59(3):211–222. Doi: 10.1002/em.22163.

2. Albayrak L, Türksoy VA, Khalilov R, et al. (2023) ‘Investigation of heavy metal exposure and trace element levels in acute exacerbatıon of COPD’, Journal of King Saud University – Science, 35(1), p. 102422. 10.1016/j.jksus.2022.102422.

3. Berge SR and Skyberg K (2003) ‘Radiographic evidence of pulmonary fibrosis and possible etiologic factors at a nickel refinery in Norway’, Journal of Environmental Monitoring, 5(4), pp. 681–688. 10.1039/B209623B.

4. Campen MJ, Nolan JP, Schladweiler MC, et al. (2001) ‘Cardiovascular and thermoregulatory effects of inhaled PM-associated transition metals: a potential interaction between nickel and vanadium sulfate’, Toxicological Sciences: An Official Journal of the Society of Toxicology, 64(2), pp. 243–252. 10.1093/toxsci/64.2.243.

5. Cao S, Yin H, Li X, et al. (2024) ‘Nickel induces epithelial-mesenchymal transition in pulmonary fibrosis in mice via activation of the oxidative stress-mediated TGF-β1/Smad signaling pathway’, Environmental Toxicology, 39(6), pp. 3597–3611. 10.1002/tox.24229.

6. Capasso L, Camatini M, and Gualtieri M (2014) ‘Nickel oxide nanoparticles induce inflammation and genotoxic effect in lung epithelial cells’, Toxicology Letters, 226(1), pp. 28–34. 10.1016/j.toxlet.2014.01.040.

7. Chandrasekar B, Colston JT, de la Rosa SD, et al. (2003) ‘TNF-alpha and H2O2 induce IL-18 and IL-18R beta expression in cardiomyocytes via NF-kappa B activation.’, Biochem Biophys Res Commun, 303(4):1152–8. doi: 10.1016/s0006-291x(03)00496-0.

8. Cheek J, Fox SS, Lehmler H-J, et al. (2024) ‘Environmental Nickel Exposure and Cardiovascular Disease in a Nationally Representative Sample of U.S. Adults’, Exposure and Health, 16(2), pp. 607–615. 10.1007/s12403-023-00579-4.

9. Chen C-Y, Lin T-K, Chang Y-C, et al. (2010) ‘Nickel(II)-induced oxidative stress, apoptosis, G2/M arrest, and genotoxicity in normal rat kidney cells’, Journal of Toxicology and Environmental Health. Part A, 73(8), pp. 529–539. 10.1080/15287390903421250.

10. Di Bucchianico S, Cappellini F, Le Bihanic F, et al. (2017) ‘Genotoxicity of TiO2 nanoparticles assessed by mini-gel comet assay and micronucleus scoring with flow cytometry’, Mutagenesis, 32(1), pp. 127–137. 10.1093/mutage/gew030.

11. Di Bucchianico S, Gliga AR, Åkerlund E, et al. (2018) ‘Calcium-dependent cyto- and genotoxicity of nickel metal and nickel oxide nanoparticles in human lung cells.’, Part Fibre Toxicol, 15(1):32. Doi: 10.1186/s12989-018-0268-y.

12. Du A, Feng S, Zhou X, et al. (2025) ‘Effect of NiCl2 Intake Through Respiratory Tract on Antioxidant Capacity, Lung, and Trace Element Content in Mice’, Biological Trace Element Research, 203(12), pp. 6144–6155. 10.1007/s12011-025-04630-0.

13. Elder A, Oberdörster G. (2006) ‘Translocation and effects of ultrafine particles outside of the lung.’, Clin Occup Environ Med, 5(4):785–96. Doi: 10.1016/j.coem.2006.07.003.

14. European Food Safety Authority (EFSA) (2020) ‘Update of the risk assessment of nickel in food and drinking water’, EFSA Journal, 18(11), p. e06268. 10.2903/j.efsa.2020.6268

15. European Scientific Committee on Occupational Exposure Limits (SCOEL) (2011) Recommendation from the Scientific Committee on Occupational Exposure Limits for nickel and inorganic nickel compounds. SCOEL/SUM/85. Luxembourg: European Commission.

16. Fidan EB, Bali EB, and Apaydin FG (2024) ‘Comparative study of nickel oxide and nickel oxide nanoparticles on oxidative damage, apoptosis and histopathological alterations in rat lung tissues’, Journal of Trace Elements in Medicine and Biology, 83, p. 127379. 10.1016/j.jtemb.2023.127379.

17. Forti E, Salovaara S, Cetin Y, et al. (2011) ‘*In vitro* evaluation of the toxicity induced by nickel soluble and particulate forms in human airway epithelial cells’, Toxicology in Vitro, 25(2), pp. 454–461. 10.1016/j.tiv.2010.11.013.

18. Freitas M, Gomes A, Porto G, et al. (2010) ‘Nickel induces oxidative burst, NF-κB activation and interleukin-8 production in human neutrophils’, Journal of biological inorganic chemistry: JBIC: a publication of the Society of Biological Inorganic Chemistry, 15(8), pp. 1275–1283. 10.1007/s00775-010-0685-3.

19. Garcés M, Marchini T, Cáceres L, et al. (2021) ‘Oxidative metabolism in the cardiorespiratory system after an acute exposure to nickel-doped nanoparticles in mice’, Toxicology, 464, p. 153020. 10.1016/j.tox.2021.153020.

20. Genchi G, Carocci A, Lauria G, et al. (2020) ‘Nickel: Human Health and Environmental Toxicology’, International Journal of Environmental Research and Public Health, 17(3):679. 10.3390/ijerph17030679.

21. Gliga AR, Di Bucchianico S, Åkerlund E, Karlsson HL. (2020) ‘Transcriptome Profiling and Toxicity Following Long-Term, Low Dose Exposure of Human Lung Cells to Ni and NiO Nanoparticles—Comparison with NiCl2.’, Nanomaterials, 10(4):649. 10.3390/nano10040649

22. Gorr MW, Youtz DJ, Eichenseer CM, et al. (2015) ‘In vitro particulate matter exposure causes direct and lung-mediated indirect effects on cardiomyocyte function’, American Journal of Physiology – Heart and Circulatory Physiology, 309(1), pp. H53–H62. 10.1152/ajpheart.00162.2015.

23. Gül U, Cakmak SK, Olcay I, et al. (2007) ‘Nickel sensitivity in asthma patients’, The Journal of Asthma: Official Journal of the Association for the Care of Asthma, 44(5), pp. 383–384. 10.1080/02770900701364213.

24. Hammond ME, Lapointe GR, Feucht PH, et al. (1995) ‘IL-8 induces neutrophil chemotaxis predominantly via type I IL-8 receptors’, Journal of Immunology, 155(3), pp. 1428–1433.

25. Hartung T (2024) ‘The (misleading) role of animal models in drug development’, Frontiers in Drug Discovery, 4. 10.3389/fddsv.2024.1355044.

26. Hobai IA, Hancox JC, and Levi AJ (2000) ‘Inhibition by nickel of the L-type Ca channel in guinea pig ventricular myocytes and effect of internal cAMP’, American Journal of Physiology-Heart and Circulatory Physiology, 279(2), pp. H692–H701. 10.1152/ajpheart.2000.279.2.H692.

27. Honda A, Tsuji K, Matsuda Y, et al. (2015) ‘Effects of air pollution-related heavy metals on the viability and inflammatory responses of human airway epithelial cells’, International Journal of Toxicology, 34(2), pp. 195–203. 10.1177/1091581815575757.

28. Hong CS, Oh SH, Lee HC, et al. (1986) ‘Occupational Asthma Caused by Nickel and Zinc’, The Korean Journal of Internal Medicine, 1(2), pp. 259–262. 10.3904/kjim.1986.1.2.259.

29. International Agency for Research on Cancer (IARC) (1990) Chromium, Nickel and Welding. IARC Monographs on the Evaluation of Carcinogenic Risks to Humans, Volume 49. Lyon, France: International Agency for Research on Cancer.

30. Izumi-Nakaseko H, Hagiwara-Nagasawa M, Naito AT, et al. (2018) ‘Application of human induced pluripotent stem cell-derived cardiomyocytes sheets with microelectrode array system to estimate antiarrhythmic properties of multi-ion channel blockers’, Journal of Pharmacological Sciences, 137(4), pp. 372–378. 10.1016/j.jphs.2018.07.011.

31. Jose CC, Jagannathan L, Tanwar VS, et al. (2018) ‘Nickel exposure induces persistent mesenchymal phenotype in human lung epithelial cells through epigenetic activation of ZEB1’, Molecular carcinogenesis, 57(6), pp. 794–806. 10.1002/mc.22802.

32. Jose CC, Wang Z, Tanwar VS, et al. (2019) ‘Nickel-induced transcriptional changes persist post exposure through epigenetic reprogramming’, Epigenetics & Chromatin, 12(1), p. 75. 10.1186/s13072-019-0324-3.

33. Jose CC, Xu B, Jagannathan L, et al. (2014) ‘Epigenetic dysregulation by nickel through repressive chromatin domain disruption’, Proceedings of the National Academy of Sciences, 111(40), pp. 14631–14636. 10.1073/pnas.1406923111.

34. Kang GS, Gillespie PA, Gunnison A, et al. (2011) ‘Long-Term Inhalation Exposure to Nickel Nanoparticles Exacerbated Atherosclerosis in a Susceptible Mouse Model’, Environmental Health Perspectives, 119(2), pp. 176–181. 10.1289/ehp.1002508.

35. Kim Y-J, Tamadon A, Kim Y-Y, et al. (2021) ‘Epigenetic Regulation of Cardiomyocyte Differentiation from Embryonic and Induced Pluripotent Stem Cells’, International Journal of Molecular Sciences, 22(16), p. 8599. 10.3390/ijms22168599.

36. Lacroix G, Koch W, Ritter D, et al. (2018) ‘Air–Liquid Interface In Vitro Models for Respiratory Toxicology Research: Consensus Workshop and Recommendations’, Applied in Vitro Toxicology, 4(2), pp. 91–106. 10.1089/aivt.2017.0034.

37. Latvala S, Hedberg J, Di Bucchianico S, et al. (2016) ‘Nickel Release, ROS Generation and Toxicity of Ni and NiO Micro- and Nanoparticles’, PloS ONE, 11(7), p. e0159684. 10.1371/journal.pone.0159684.

38. Lenz A-G, Karg E, Brendel E, et al. (2013) ‘Inflammatory and Oxidative Stress Responses of an Alveolar Epithelial Cell Line to Airborne Zinc Oxide Nanoparticles at the Air-Liquid Interface: A Comparison with Conventional, Submerged Cell-Culture Conditions’, BioMed Research International, 2013(1), p. 652632. 10.1155/2013/652632.

39. Lippmann M, Ito K, Hwang J-S, et al. (2006) ‘Cardiovascular Effects of Nickel in Ambient Air’, Environmental Health Perspectives [Preprint]. 10.1289/ehp.9150.

40. Lou S, Zhong L, Yang X, et al. (2013) ‘Efficacy of all-*trans* retinoid acid in preventing nickel induced cardiotoxicity in myocardial cells of rats’, Food and Chemical Toxicology, 51, pp. 251–258. 10.1016/j.fct.2012.09.007.

41. Lü X, Bao X, Huang Y, et al. (2009) ‘Mechanisms of cytotoxicity of nickel ions based on gene expression profiles’, Biomaterials, 30(2), pp. 141–148. 10.1016/j.biomaterials.2008.09.011.

42. Mau R, Kussauer S, Matzmohr U, et al. (2021) ‘Customised micro-electrode array (MEA) test setup featuring a silicone-casted overlay with two chambers for separated cell seedings’, Current Directions in Biomedical Engineering, 7(2), pp. 311–314. 10.1515/cdbme-2021-2079.

43. Miller JM, Mardhekar NM, Pretorius D, et al. (2020) ‘DNA damage-free iPS cells exhibit potential to yield competent cardiomyocytes’, American Journal of Physiology. Heart and Circulatory Physiology, 318(4), pp. H801–H815. 10.1152/ajpheart.00658.2019.

44. Mo Y, Zhang Y, and Zhang Q (2024) ‘The pulmonary effects of nickel-containing nanoparticles: Cytotoxicity, genotoxicity, carcinogenicity, and their underlying mechanisms’, Environmental science. Nano, 11(5), pp. 1817–1846. 10.1039/d3en00929g.

45. Nakane H (2012) ‘Translocation of particles deposited in the respiratory system: a systematic review and statistical analysis’, Environmental Health and Preventive Medicine, 17(4), pp. 263–274. 10.1007/s12199-011-0252-8.

46. Ng WH, Johnston EK, Tan JJ, et al. (2022) ‘Recapitulating human cardio-pulmonary co-development using simultaneous multilineage differentiation of pluripotent stem cells’, eLife. Edited by P.W. Noble and S.X. Huang, 11, p. e67872. 10.7554/eLife.67872.

47. Ouyang W, Zhang D, Li J, et al. (2009) ‘Soluble and insoluble nickel compounds exert a differential inhibitory effect on cell growth through IKKα-dependent cyclin D1 down-regulation’, Journal of cellular physiology, 218(1), pp. 205–214. 10.1002/jcp.21590.

48. Pietruska JR, Liu X, Smith A, et al. (2011) ‘Bioavailability, Intracellular Mobilization of Nickel, and HIF-1α Activation in Human Lung Epithelial Cells Exposed to Metallic Nickel and Nickel Oxide Nanoparticles’, Toxicological Sciences, 124(1), pp. 138–148. 10.1093/toxsci/kfr206.

49. Pourrier M and Fedida D (2020) ‘The Emergence of Human Induced Pluripotent Stem Cell-Derived Cardiomyocytes (hiPSC-CMs) as a Platform to Model Arrhythmogenic Diseases’, International Journal of Molecular Sciences, 21(2), p. 657. 10.3390/ijms21020657.

50. Refsvik T and Andreassen T (1995) ‘Surface binding and uptake of nickel(II) in human epithelial kidney cells: modulation by ionomycin, nicardipine and metals’, Carcinogenesis, 16(5), pp. 1107–1112. 10.1093/carcin/16.5.1107.

51. Rizwan M, Usman K, and Alsafran M (2024) ‘Ecological impacts and potential hazards of nickel on soil microbes, plants, and human health’, Chemosphere, 357, p. 142028. 10.1016/j.chemosphere.2024.142028.

52. Schaumlöffel D (2012) ‘Nickel species: Analysis and toxic effects’, Journal of Trace Elements in Medicine and Biology, 26(1), pp. 1–6. 10.1016/j.jtemb.2012.01.002.

53. Shinohara N, Zhang G, Oshima Y, et al. (2017) ‘Kinetics and dissolution of intratracheally administered nickel oxide nanomaterials in rats’, Particle and Fibre Toxicology, 14, p. 48. 10.1186/s12989-017-0229-x.

54. Stinson TJ, Jaw S, Jeffery EH, et al. (1992) ‘The relationship between nickel chloride-induced peroxidation and DNA strand breakage in rat liver’, Toxicology and Applied Pharmacology, 117(1), pp. 98–103. 10.1016/0041-008x(92)90222-e.

55. Suetomi T, Willeford A, Brand CS, et al. (2018) ‘Inflammation and NLRP3 Inflammasome Activation Initiated in Response to Pressure Overload by Ca2+/Calmodulin-Dependent Protein Kinase II δ Signaling in Cardiomyocytes Are Essential for Adverse Cardiac Remodeling.’, Circulation, 138(22):2530–2544. doi: 10.1161/CIRCULATIONAHA.118.034621.

56. Tajbakhsh J, Mortazavi F, and Gupta NK (2022) ‘DNA methylation topology differentiates between normal and malignant in cell models, resected human tissues, and exfoliated sputum cells of lung epithelium’, Frontiers in Oncology, 12. 10.3389/fonc.2022.991120.

57. Wallenborn JG, McGee JK, Schladweiler MC, et al. (2007) ‘Systemic Translocation of Particulate Matter–Associated Metals Following a Single Intratracheal Instillation in Rats’, Toxicological Sciences, 98(1), pp. 231–239. 10.1093/toxsci/kfm088.

58. Wang H, Wang T, Rui W, et al. (2022) ‘Extracellular vesicles enclosed-miR-421 suppresses air pollution (PM2.5)-induced cardiac dysfunction via ACE2 signalling’, Journal of Extracellular Vesicles, 11(5), p. e12222. 10.1002/jev2.12222.

59. Wang M, Markel TA, Meldrum DR. (2008) ‘Interleukin 18 in the heart.’, Shock, 30(1):3–10. doi: 10.1097/SHK.0b013e318160f215.

60. Wu C-H, Hsiao Y-M, Yeh K-T, et al. (2017) ‘Upregulation of microRNA-4417 and Its Target Genes Contribute to Nickel Chloride-promoted Lung Epithelial Cell Fibrogenesis and Tumorigenesis’, Scientific Reports, 7(1), p. 15320. 10.1038/s41598-017-14610-7.

61. Yu J, Zhang H, Wu F, et al. (2006) ‘Insulin improves cardiomyocyte contractile function through enhancement of SERCA2a activity in simulated ischemia/reperfusion’, Acta Pharmacologica Sinica, 27(7), pp. 919–926. 10.1111/j.1745-7254.2006.00388.x.

62. Zambelli B, Uversky VN, and Ciurli S (2016) ‘Nickel impact on human health: An intrinsic disorder perspective’, Biochimica et Biophysica Acta (BBA) - Proteins and Proteomics, 1864(12), pp. 1714–1731. 10.1016/j.bbapap.2016.09.008.

63. Zhang N, Chen M, Li J, et al. (2019) ‘Metal nickel exposure increase the risk of congenital heart defects occurrence in offspring: A case-control study in China’, Medicine, 98(18), p. e15352. 10.1097/MD.0000000000015352.

64. Zheng J, Wang J, Li K, et al. (2023) ‘LncRNA AP000487.1 regulates PRKCB DNA methylation-mediated TLR4/MyD88/NF-κB pathway in Nano NiO-induced collagen formation in BEAS-2B cells’, Environmental Toxicology, 38(11), pp. 2783–2796. 10.1002/tox.23918.

65. Zheng J, Wang J, Qin X, et al. (2022) ‘LncRNA HOTAIRM1 Involved in Nano NiO-Induced Pulmonary Fibrosis via Regulating PRKCB DNA Methylation-Mediated JNK/c-Jun Pathway’, Toxicological Sciences, 190(1), pp. 64–78. 10.1093/toxsci/kfac092.

66. Zhou X, Liao J, Lei Z, et al. (2025) ‘Nickel-based nanomaterials: a comprehensive analysis of risk assessment, toxicity mechanisms, and future strategies for health risk prevention’, Journal of Nanobiotechnology, 23, p. 211. 10.1186/s12951-025-03248-7.

67. Zimmermann EJ, Candeias J, Gawlitta N, et al. (2023) ‘Biological impact of sequential exposures to allergens and ultrafine particle-rich combustion aerosol on human bronchial epithelial BEAS-2B cells at the air liquid interface’, Journal of Applied Toxicology, 43(8), pp. 1225–1241. 10.1002/jat.4458.

